# Whole-brain neural substrates of behavioral variability in the larval zebrafish

**DOI:** 10.1101/2024.03.03.583208

**Authors:** Jason Manley, Alipasha Vaziri

## Abstract

Animals engaged in naturalistic behavior can exhibit a large degree of behavioral variability even under sensory invariant conditions. Such behavioral variability can include not only variations of the same behavior, but also variability across qualitatively different behaviors driven by divergent cognitive states, such as fight-or-flight decisions. However, the neural circuit mechanisms that generate such divergent behaviors across trials are not well understood. To investigate this question, here we studied the visual-evoked responses of larval zebrafish to moving objects of various sizes, which we found exhibited highly variable and divergent responses across repetitions of the same stimulus. Given that the neuronal circuits underlying such behaviors span sensory, motor, and other brain areas, we built a novel Fourier light field microscope which enables high-resolution, volumetric imaging across the larval zebrafish during behavior. This enabled us to screen for neural loci which exhibited activity patterns correlated with behavioral variability. We found that despite the highly variable activity of neurons, visual stimuli were robustly encoded at the population level, and the visual-encoding dimensions of neural activity did not explain behavioral variability. This robustness despite apparent neuronal variability was due to the multi-dimensional geometry of the neuronal population dynamics: almost all neural dimensions that were variable across individual trials, i.e. the “noise” modes, were nearly orthogonal to those encoding for sensory information. Investigating this neuronal variability further, we identified two sparsely-distributed, brain-wide neuronal populations whose pre-motor activity predicted whether the larva would respond to a stimulus and, if so, which direction it would turn on a single-trial level. These populations predicted single-trial behavior seconds before stimulus onset, indicating they encoded time-varying internal states modulating behavior, perhaps organizing behavior over longer timescales or enabling flexible behavior routines. Our results provide the first brain-wide evidence that sensory, motor, and internal variables are represented in a highly mixed fashion across widely distributed neuronal populations and demonstrate that de-mixing each of these components at the neuronal population level is critical to understanding the mechanisms underlying the brain’s remarkable flexibility and robustness.

## INTRODUCTION

Animals are not deterministic input-output machines, but instead display highly flexible behavioral responses even under sensory invariant conditions. A holistic understanding of the neuronal mechanisms underlying decision making requires explaining such variability in behavior at the single trial level. However, there are many putative sources of trial-to-trial neuronal variability. In particular, it is unclear whether such variability results from deterministic sources, such as internal states and long-timescale dynamics, or stochasticity. Individual neurons exhibit non-negligible noise from a variety of sources, including electrical noise due to ion channel fluctuations, synaptic noise due to biochemical processes, jitter in the timing of action potentials, and “digitization” noise due to the all-or-nothing nature of the action potential (Faisal et al., 2008).

However, trial-to-trial variability in neuronal firing is generally not independent across neurons, but instead highly correlated over repeated stimulus presentations within certain groups of neurons (Kohn et al., 2015; Shadlen and Newsome, 1998; Zohary et al., 1994), suggesting that variability in neural circuits may not be dominated by the stochasticity of individual neurons. Such structured covariation is referred to as “noise correlations” (Cohen and Kohn, 2011) because they are unrelated to the often experimenter-controlled external variable that is simultaneously encoded within the neuronal population. While certain structures of noise correlations are known to limit the information encoding capacity of the neuronal population (Moreno-Bote et al., 2014; Bartolo et al., 2020; Rumyantsev et al., 2020), there is little consensus about both their origin (Kanitscheider et al., 2015; Zylberberg et al., 2016) and behavioral impact (Zohary et al., 1994; Huang and Lisberger, 2009). In particular, there are opposing reports in different species and under different task conditions as to whether such inter-neuronal correlations can improve (Cafaro and Rieke, 2010; Zylberberg et al., 2016; Valente et al., 2021) or interfere (Cohen and Maunsell, 2009; Ruda et al., 2020; Kafashan et al., 2021) with the neural computations underlying decision making. As such, a general framework capable of reconciling such widely varying reports and explaining the functional impact of noise correlations on neural computation and behavior is currently missing.

Further, few experimental studies have linked noise correlations in sensory regions to pre-motor dynamics and ultimately behavior (Valente et al., 2021). Instead, most reports have mainly focused on the scale and structure of such covariations across sensory neurons in anesthetized animals or without regard to the animal’s behavioral state. However, recent studies have shown that many species encode the animal’s own movements and decisions in a highly distributed fashion, including across early sensory regions (Kauvar et al., 2020; Musall et al., 2019; Stringer et al., 2019; Steinmetz et al., 2019). These brain-wide, motor-related fluctuations must then drive at least some of the observed neuronal covariation across stimulus presentations. Thus, noise correlations are not merely stochastic “noise”, but instead may encode important variables that are stimulus-independent and perhaps related to the animal’s proprioception and context, such as long-timescale behavioral and internal state dynamics.

Such highly overlapping representations of sensory information and additional contextual variables may underlie the brain’s remarkable ability to flexibly control behavior and select the most appropriate action at any given moment based on the animal’s goals and internal state. As such, naturalistic decision making is a function of not just incoming sensory stimuli, but also internal variables such as the animal’s motivations and recent experience or learned behaviors. For example, foraging has been shown across species to be modulated by hunger (Filosa et al., 2016), long-timescale exploration versus exploitation states (Flavell et al., 2013; Marques et al., 2020), and other hidden internal variables (Sternson, 2020; Torigoe et al., 2021; Flavell et al., 2022). This evidence suggests that answering longstanding questions regarding the neuronal mechanisms underlying naturalistic decision making will require understanding the intersection of numerous neural circuits distributed throughout the vertebrate brain.

Thus, probing the structure of the neuronal mechanisms underlying behavioral variability requires high spatiotemporal resolution recording of brain-wide neuronal dynamics and behavior across many trials. While it is often necessary to average across trials to deal with both the inherent variable firing of neurons and experimental measurement noise, such trial averaging inherently precludes any investigation of the link between neural and behavioral variability on the trial-by-trial level. Instead, methods that utilize the entire neuronal population dynamics have shown that pooling information across neurons can allow for successful decoding of information that is not possible from individual neurons. This is because simultaneous multi-neuron recordings can leverage statistical power across neurons to capture the most salient aspects of population dynamics within single trials. Such population-level approaches have revolutionized the fields of motor planning and decision making, providing highly robust brain-machine interfaces (Kao et al., 2015; Pandarinath et al., 2018) that combine information from at least a few recording sites and making it possible to predict decisions (Wei et al., 2019; Lin et al., 2020), reaction time (Afshar et al., 2011), and additional behavioral states (Kaufman et al., 2015; Marques et al., 2020) at the single trial level. Recently, optical imaging techniques (Weisenburger and Vaziri, 2018; Kim and Schnitzer, 2022; Machado et al., 2022) have dramatically increased the attainable number of simultaneously recorded neurons within increasingly larger brain volumes at high speed and resolution. Thus, large-scale neural recording provides a promising approach to both screening for potential sources of variability across the brain and identifying robust population-level dynamics at the level of single trials. However, due to the difficulty of obtaining whole-brain optical access in many vertebrates, previous studies have not been able to quantify brain-wide neuronal noise correlations and link such noise modes to variable decisions at the single trial level.

In this study, we sought to identify the neural loci which drive or encode information related to behavioral variability across trials. To do so, we performed brain-wide recording in larval zebrafish engaged in ethologically relevant visually-evoked behaviors to identify neuronal populations that were predictive of behavioral variability under sensory invariant conditions. While it is well-known that larval zebrafish on average exhibit a target-directed prey capture response to small visual stimuli and a target-avoidance escape response to large visual stimuli, we demonstrated that these ethological, oppositely valenced, and highly divergent responses exhibited significant variability across trials of identical stimulus presentations. Turning first to visually-evoked neural populations, we found that despite trial-to-trial variability in the responses of individual units, visual information was reliably encoded at the population level across trials of each stimulus. As such, the visual-encoding neuronal activity patterns did not account for the larvae’s variable behavioral responses across trials.

A key feature of our system was that it allowed investigating the geometry of trial-to-trial variability of neuronal dynamics across the brain, by simultaneous observation of brain-wide neural variability during a visual decision-making behavior. Thus, we were able to identify brain-wide modes of neural variability which were related to the larva’s motor actions. In particular, we identified highly distributed neuronal populations whose pre-motor activity predicted the larvae’s responsiveness and turn direction on a single trial level, indicating that behavioral variability is not represented in particular anatomical neural loci. We found that this predictability exhibited two dominant timescales: a longer-timescale and pre-stimulus turn bias that was not time-locked to the stimulus presentations or motor timing; and a rapid increase in predictability and ramping activity about one to two seconds before movement initiation. Consistent with results from previous studies, we speculate that the former, longer-timescale bias is related to a circuitry that has evolved to optimize an exploration versus exploitation behavior in foraging while the latter, short-timescale ramping activity likely drives the downstream motor circuitry to execute the actual selected action in each trial. Our data suggest that behavioral variability in response to repetitions of the same sensory stimulus may not be driven by a single brain region. Rather, it is more likely generated by a combination of factors encoded by neurons throughout the brain, including a time-varying and internal turn direction bias, in addition to the well-studied visuomotor transformations.

## RESULTS

### Fourier Light Field Microscopy enables high-speed imaging of neuronal population dynamics

In order to investigate the brain-wide neural correlates of behavioral variability, we designed and built a Fourier light field microscope (fLFM) that was optimized for high-resolution recording of neuronal activity in the larval zebrafish. Conventional light field microscopy (cLFM) provides an elegant solution to volumetric snapshot imaging by placing a microlens array (MLA) in the image plane to encode both the position and angle of incidence of the light rays onto a camera sensor (Levoy et al., 2006). Combined with 3D deconvolution algorithms for computational reconstruction of the recorded volume (Agard, 1984; Broxton et al., 2013; Prevedel et al., 2014), LFM enables imaging of an entire volume without requiring any scanning. Due to its high speed and volumetric capabilities, cLFM has been widely applied to image the dynamics of *in vivo* biological systems, particularly neuronal dynamics (Prevedel et al., 2014; Aimon et al., 2019; Lin et al., 2020; Quicke et al., 2020; Yoon et al., 2020). Further, additional computational strategies have been devised to image deep into scattering tissue, such as the mouse brain (Nöbauer et al., 2017; Skocek et al., 2018; Nöbauer et al., 2023). However, a primary limitation of cLFM is a dramatic drop in resolution near the native image plane (NIP) because its spatio-angular sampling is uneven across depth and highly redundant near the NIP.

This low resolution at the center of the sample can only be partially overcome using techniques such as wavefront coding (Cohen et al., 2014). In contrast, fLFM has been shown to provide high-resolution imaging across a two- to three-fold extended depth by processing the light field information through the Fourier domain (Llavador et al., 2016; Scrofani et al., 2017; Cong et al., 2017; Guo et al., 2019; Liu et al., 2020); i.e., placing the MLA conjugate to the back focal plane of the objective. In this configuration, the MLA segments the wavefront by transmitting the various spatial frequencies (which correspond to angular information) into images on different regions of the camera, providing a well-aliased sampling of the spatio-angular information without significant redundancy near the NIP.

We simultaneously optimized our design for simplicity and cost-effectiveness by using exclusively off-the-shelf elements, as opposed to previous implementations of fLFM which relied on custom optical components. Our design (Figure 1A, see Methods for details) enables imaging of an approximately 750 × 750 × 280 μm^3^ volume, corresponding to roughly the whole brain of a 7-day-post-fertilization (dpf) larval zebrafish. Here, “whole-brain” refers to the anatomical extent of the imaging volume; it does not imply uniform labeling, neuronal sampling, or cellular separability across all regions. 3D information is captured in the raw images (Figure 1B) because each lenslet in the MLA segments a region of the Fourier plane and thus forms an image of the sample from a unique perspective. Experimentally, the point spread function (PSF) full width at half maximum (FWHM) of our system measured 3.3 ± 0.21 μm (95% confidence interval) laterally and 5.4 ± 0.4 μm axially (Figure 1C), consistent with theoretical estimates of 3.1 μm and 4.7 μm, respectively. This represents a substantial improvement in resolution compared to our previous cLFM, which had a PSF FWHM of 3.4 μm laterally and 11.3 μm axially (Prevedel et al., 2014). Additionally, the fLFM has a much more consistent resolution as a function of depth as compared to the cLFM (Figure S1A), which exhibits a dramatic drop in resolution near the NIP. These measurements provide a standardized characterization of fLFM optical performance under controlled calibration conditions. Previous work combining *in vivo* backscatter measurements with experimentally constrained modeling at a comparable visible wavelength found relatively modest focal broadening over the first 100 µm of larval zebrafish brain, including within nucleus-rich regions (Favre-Bulle et al., 2015). Nevertheless, cellular separability in vivo may vary with imaging depth, labeling density, signal quality, and reconstruction.

**Figure 1:**
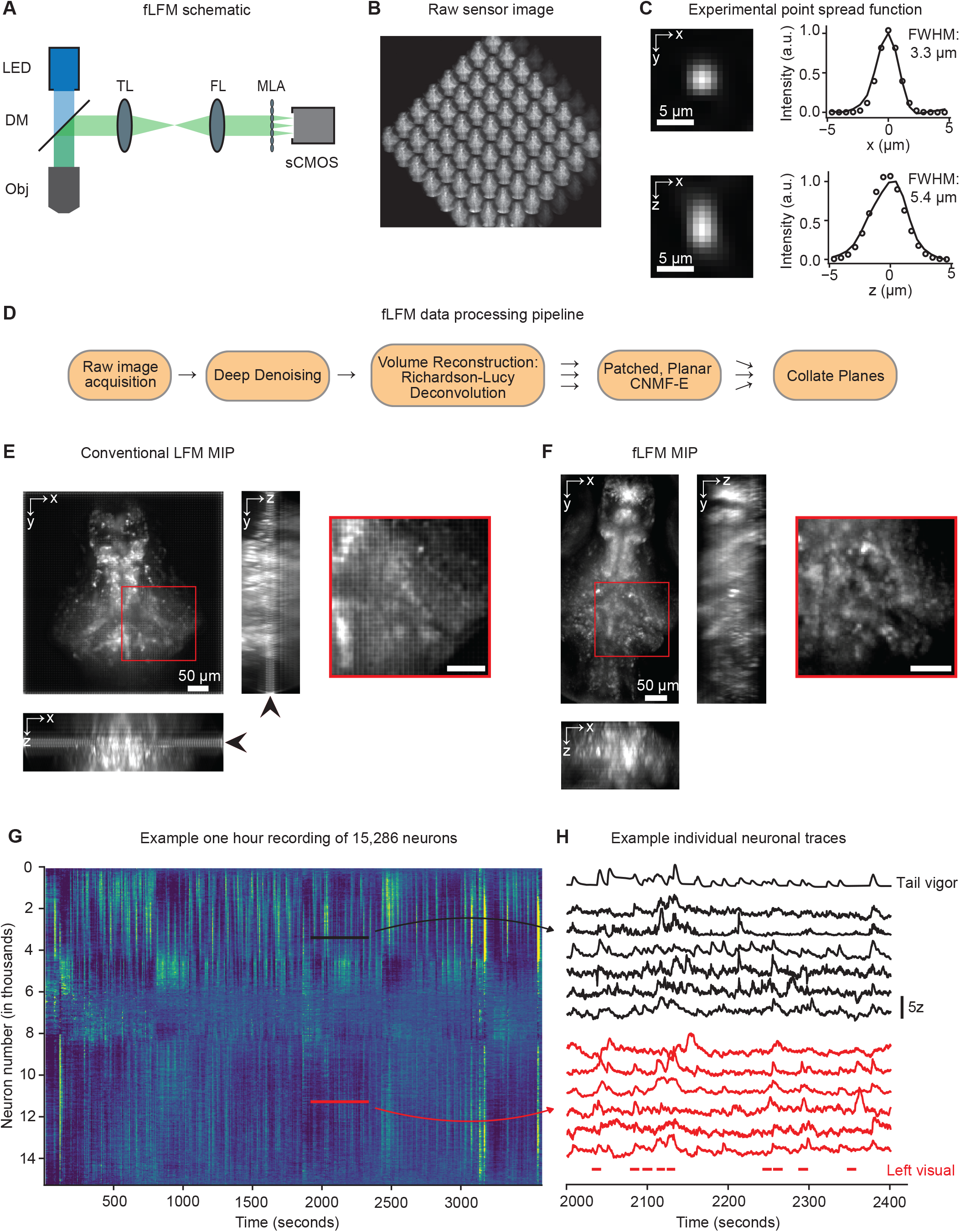
Fourier Light Field Microscopy (fLFM) provides a simple and cost-effective method for high-speed volumetric imaging across the larval zebrafish brain during behavior. A. Schematic of the fLFM system. The sample is illuminated with a 470 nm LED through a 20×/1.0-NA imaging objective (Obj). The fluorescence is sent to the imaging path via a dichroic mirror (DM). An Olympus-style f=180mm tube lens (TL) and f=180mm Fourier lens (FL) are used to conjugate the back focal plane of the objective onto a microlens array (MLA). An sCMOS sensor is positioned at the focal plane of the MLA to capture the raw fLFM images. **B.** An example raw sensor image. Each lenslet in the 8×8 array forms an image of the sample from a slightly different perspective, allowing for reconstruction of the 3D volume. **C.** Experimental measurement of the point spread function (PSF). Left: the x-y (top) and x-z (bottom) profiles of a reconstructed image of a 1 μm fluorescent bead embedded in 2.5% agarose. Right: corresponding cross sections of the PSF (points). A Gaussian profile (lines) was fit to these data to measure the full width at half maximum (FWHM), which was 3.3 μm laterally and 5.4 μm axially. **D.** Schematic of our fLFM data processing pipeline (see Methods for detailed description). **E.** A maximum intensity projection (MIP) of a conventional LFM (cLFM) larval zebrafish recording resulting in strong grid-like artifacts and low resolution near the native image plane (black arrows). A reconstructed volume of 750 × 750 × 200 μm^3^ is shown. **F.** An MIP of an fLFM recording. fLFM exhibits higher axial resolution and does not contain artifacts at the native image plane. A reconstructed volume of 750 × 375 × 250 μm^3^ is shown. **G.** A heatmap of extracted neuronal activity from an example one hour recording of 15,286 neuronal ROIs. Neurons are sorted using the rastermap algorithm (Stringer et al., 2019), such that nearby ROIs exhibit similar temporal activity patterns. Multiple distinct activity patterns are seen, including strong activity during two sets of drifting grating-induced optomotor response trials (black arrows). **H.** Example neurons tuned to tail movements and visual stimuli. In black are six example neuron traces from the designated region in panel G, which exhibited correlations with the GCaMP-kernel-convolved tail vigor (top trace, see Methods for details). In red are six example neuron traces from the designated region in panel G, which exhibited correlations with the visual stimuli presented in the left visual field (denoted by the red lines at the bottom of the plot).

Next, we developed a custom pipeline (Figure 1D) for 3D reconstruction and neuronal segmentation to extract the dynamics of neurons across the field of view. This consisted of three main steps: denoising, 3D reconstruction, and neuronal segmentation. As one of the key limitations of LFM is signal-to-noise ratio (SNR) due to the distribution of emitted photons onto a sensor array, we first denoised the raw sensor images. To do so we trained a DeepInterpolation (Lecoq et al., 2021) deep neural network model, which has been shown to increase the SNR of the resulting neuronal timeseries by removing shot noise that is independent across consecutive frames (Figure S1B, see Methods for details). Next, the full 3D volume was reconstructed from denoised sensor images using Richardson-Lucy deconvolution (Agard, 1984; Broxton et al., 2013; Prevedel et al., 2014) and an experimentally measured PSF. In addition to enabling high-resolution imaging across an extended depth of field, a key advantage of fLFM is that the reconstruction of the volume images can be performed approximately 100 times faster than in cLFM, due to fLFM’s shift-invariant point spread function. Finally, to identify neuronal regions of interest (ROIs) and their dynamics within the reconstructed volume, we utilized CNMF-E (Zhou et al., 2018), a constrained matrix factorization approach to extract *in vivo* calcium signals from one-photon imaging data. The CNMF-E algorithm was applied to each plane in parallel, after which the neuronal ROIs from each plane were collated and duplicate ROIs across planes were merged (Figure S1C). Finally, the effect of photobleaching was corrected according to a biexponential fit to the mean activity across ROIs (Figure S1D).

To validate our setup, we imaged brain-wide dynamics from head-immobilized larval zebrafish expressing nuclear-localized GCaMP6s (NL-GCaMP6s) pan-neuronally (Vladimirov et al., 2014) while monitoring the larva’s tail movements using a high-speed camera and presenting visual stimuli, which as we will discuss below consisted of single dots of 3-5 different sizes moving from the center toward the left or right visual field. Across our 1-2 hour recordings, we identified 16,524 ± 3,942 neuronal ROIs per animal (mean ± 95% confidence interval; range: 6,425 to 35,928) distributed across all major brain divisions represented in the recording volume. Their aggregate distribution broadly resembled the Z-Brain reference image (Randlett et al., 2015) labeled with nuclear-localized red fluorescent protein (Figure S1E). Thus, the combination of fLFM and the improvement in our reconstruction pipeline enabled the detection of significantly more neuronal ROIs than the ∼5,000 ROIs reported in our previous cLFM realizations (Prevedel et al., 2014; Lin et al., 2020). The performance improvement over cLFM could also be visualized by inspecting the maximum intensity projection (MIP) of a zebrafish recording in each modality. While cLFM (Figure 1E) exhibits lower resolution and grid-like artifacts near the native image plane (black arrows) due to redundant sampling near the NIP, fLFM’s well-aliased sampling of spatio-angular information substantially improves resolution throughout the volume (Figure 1F). Although the larval zebrafish brain is relatively transparent and only weakly scattering, which should mitigate depth-dependent degradation relative to more opaque vertebrate preparations, *in vivo* cellular separability at depth can still vary with reporter brightness, neuronal density, and residual scattering.

Within these observed large-scale dynamics, we found a diversity of neuronal activity patterns (Figure 1G). For example, we identified neuronal ROIs tuned to the vigor of the larva’s tail movements (see Methods for details) and those tuned to the presentation of visual stimuli as described below (Figure 1H), providing a proof of principle that fLFM enables brain-wide imaging of the relevant neuronal population activity encoding sensory inputs and behavior.

### Larval zebrafish exhibit highly variable motor responses to visual stimuli

fLFM therefore provides a promising approach for screening for regions across the brain which exhibited covariations related to variability in behavior and naturalistic decision making. We thus set out to investigate the neuronal basis of trial-to-trial variability in action selection during two different ethologically-relevant behaviors which are known to be generated by brain-wide sensorimotor circuitries (Chen et al., 2018). In the larval zebrafish, several visually-evoked behaviors such as prey capture (Borla et al., 2002; Patterson et al., 2013) and escape response (Temizer et al., 2015) have been widely studied. In particular, varying a single stimulus parameter, the size of a visual object, can elicit dramatically distinct average behavioral responses (Barker and Baier, 2015), ranging from target-directed responses (e.g., prey capture) to small stimuli, to target-avoidance behaviors (e.g., escape response) to large stimuli. These distinct behaviors can be evoked in the laboratory using simple visual stimuli and in a head-fixed imaging preparation (Bianco et al., 2011; Semmelhack et al., 2014; Temizer et al., 2015; Barker and Baier, 2015; Bianco and Engert, 2015; Dunn et al., 2016a; Filosa et al., 2016; Förster et al., 2020; Oldfield et al., 2020). As such, the neural circuitry underlying each of the individual behaviors is well understood (Portugues and Engert, 2009; Bollmann, 2019) on the trial-averaged level. However, importantly these behaviors are not reflexive. They involve the processing of information across multiple brain regions and are subjected to neuromodulatory effects and longer time scale changes of internal states (Filosa et al., 2016; Marques et al., 2020). As such, the larvae do not deterministically perform the same actions over multiple sensory invariant trials. Even under highly optimized and invariant conditions, prey capture is only observed in up to 30% of trials (Bianco et al., 2011), whereas escape response is more consistently observed in 80% of trials (Temizer et al., 2015). However, the neuronal populations and the underlying circuitry driving such variability in responsiveness and action selection in either of these paradigms have not been identified, yet these ethologically highly divergent behaviors can be elicited by tuning a single parameter. We thus hypothesized that modulating the size of the visual object should reveal a regime of behavioral variability between these highly divergent, naturalistic behaviors, providing a useful means and context to study questions such as how behavioral strategies may switch across trials or how neural and behavioral variability changes as a function of the ambiguity of the presented stimulus.

Thus, we first sought to characterize the trial-to-trial variability of the larvae’s behavioral responses to repetitions of simple visual stimuli of various sizes. In a freely swimming context (Figure 2A), we displayed moving dots of various sizes ranging from 1 to 40 visual degrees (see Methods for details) on a screen positioned below the larvae. Analyzing these behavioral recordings for different dot sizes, we indeed found characteristic target-avoidance bouts similar to escape response (Figure 2B) and target-directed bouts similar to prey capture response (Figure 2C; see also Video 1). For each bout of behavior in which a larva was close to a visual object, we quantified whether it approached or avoided the target using the larva’s change in distance from the object as a metric (see Methods for details). Consistent with previous work (Barker and Baier, 2015), on average the larvae exhibited target-directed behavior to small stimuli (1-5°), followed by a crossover point at which the larvae were not preferentially target-directed or target-avoiding for stimuli of 7° in size, and then a target-avoidance behavioral regime to stimuli above 10°, which elicited characteristic escape responses on average (Figure 2D). Moreover, across individual trials the larvae’s responses exhibited significant variability, yielding many trials in which the larvae either did not respond or responded in the opposite direction of the average response, as indicated by the large standard deviation across bouts for the same stimulus size (Figure 2D).

**Figure 2:**
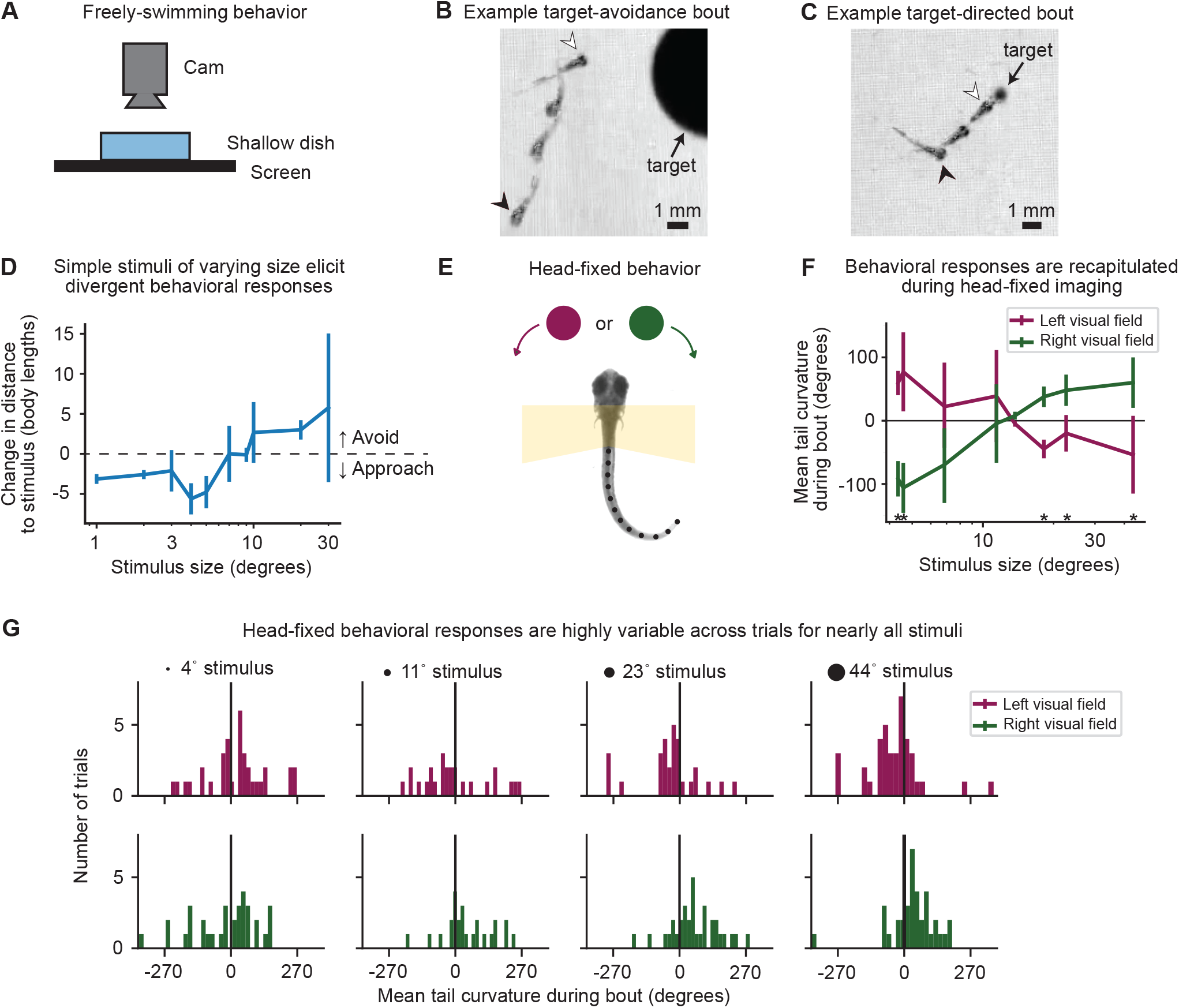
Zebrafish exhibit highly variable motor responses to visual stimuli. **A.** Schematic of the freely-behaving experimental setup. A single larva is placed in a 90 mm petri dish with a shallow (approximately 3 mm) amount of water. The dish is placed on a screen which displays visual stimuli consisting of dots of various sizes drifting in a constant direction and speed. The behavior is monitored with a camera from above. **B.** An example target-avoidance bout. A composite image taken of four frames from start (white arrow) to near the finish (black arrow) of a bout in which the larva avoided a large visual stimulus. See also Video 1. **C.** An example target-directed bout. A composite image taken of three frames from start (white arrow) to finish (black arrow) of a bout in which the larva made a directed movement toward a small visual stimulus. See also Video 1. **D.** The behavioral response of larvae to various size visual stimuli. For each size stimulus, all bouts of movement in which the larva was within six body lengths of a visual stimulus were considered. For each bout, the change in distance between the larvae and the stimulus during the movement was monitored. Thus, target-directed bouts exhibited negative values corresponding to a decrease in the distance to the stimulus, whereas target-avoidance bouts exhibited positive values. Stimuli less than 7° in size evoked target-directed responses on average, whereas stimuli greater than 10° evoked target-avoidance responses. Shown is the mean ± standard deviation of n=9 larvae. **E.** Schematic of the head-fixed experimental setup. A larva is embedded in agarose in order to keep its head fixed during brain-wide imaging. The agarose around the tail and eyes is removed, such that the tail can be tracked (indicated by black dots along the tail) and visual stimuli can be presented. Visual stimuli of various sizes are presented moving from the center of visual field to either the left or right. **F.** The behavioral response of larvae to various size visual stimuli during head fixation. Visual stimuli are presented in either the left (pink) or right (green) visual field. The directedness of tail movements is monitored by computing the mean tail curvature during a bout of movement, with positive values indicating leftward motions. Similar to freely-behaving experiments, visual stimuli of 7° or less evoked target-directed responses, whereas stimuli larger than 10° evoked target-avoidance. Shown is the mean ± standard deviation of n=10 larvae (inter- and intra-larva variability is described in Figure S2). Asterisks indicate stimuli with significant difference between presentations on the left and right visual field (p<0.05, paired t-test). **G.** Example behavioral responses to various stimuli. For each of four stimulus sizes on the left and right visual fields, a histogram of the mean tail curvature during bouts is shown for an example larva. While stimulus-evoked behavioral responses are variable in all cases, they appear the least variable in the case of largely target-avoidance bouts to large 44° stimuli (rightmost column).

Next, we confirmed that these behaviors were preserved in a head-fixed imaging preparation, in which the larvae were embedded in agarose with their eyes and tail cut free (Figure 2E), such that their variable behavioral responses to visual stimuli could be tracked while performing brain-wide neuronal recordings using our fLFM system. To do so, we presented visual stimuli as dots of a certain size moving in either the left or right visual field of the animals (see Methods for details). Visual stimuli were created using a projector that projected onto the edge of the circular 35 mm petri dish in which the larvae were embedded. Utilizing curvature of the tail to the left or the right as a metric for target-directed versus -avoidance behavior, we again identified the stimulus-size dependence of the larvae’s average behavioral response (Figures 2F, S2A) that recapitulated what was observed in freely behaving larvae. We also confirmed that the corresponding opposite tail responses were observed when the stimuli were presented on the other side of the visual field, demonstrating that even during head-fixation the larvae exhibit on average directed behaviors that depend on the stimulus size. However, at the single trial level, we observed behavioral variability to nearly all stimulus sizes (Figures 2G, S2B), even in the stimulus size regime known to optimally drive prey capture and escape response behaviors. The most widely variable responses were at 4° and 11°, corresponding to stimuli in the target-directed regime and near the crossover between target-directed and avoidance, respectively. This high degree of variability among responses to small and intermediate stimuli is consistent with previous literature estimates that even in optimized setups, the prey capture response is observed at rates less than 30% during head-fixation (Bianco et al., 2011) and any motor response at all to small visual stimuli is observed at rates up to 60% of trials in free behavior (Filosa et al., 2016). However, the neural mechanisms and circuitry underlying such trial-to-trial behavioral variability are not understood. In particular, it is unclear whether such variability within these ethologically relevant, yet divergent behaviors is driven by noise and inherent stochasticity in neuronal circuits, or represents modulation by time-varying internal states, such as motivation and hunger.

### Trial-to-trial variability in visually-evoked neurons is largely orthogonal to visual decoding dimensions

Previous studies have reported that individual neurons tuned to specific stimuli often exhibit significant variability in their responses over multiple presentations of the same stimulus (Montijn et al., 2016; Rumyantsev et al., 2020; Shadlen and Newsome, 1998; Zohary et al., 1994; Zylberberg et al., 2016). Thus, one potential mechanism underlying our observed behavioral variability could be trial-to-trial deviations in the neural encoding of visual stimuli. Alternatively, the baseline pre-stimulus state of visually-evoked neurons could be shifted by other circuit components. Given that we had observed behavioral variability across the entire studied range of stimulus sizes, we proceeded to record brain-wide dynamics with the fLFM while presenting many repetitions of 3 to 6 different stimulus sizes spanning 1° to 40° (see Methods for details). Investigating the responses of neurons across the whole brain, we indeed found that even neuronal ROIs highly tuned to a particular stimulus exhibited variability in their responses across multiple presentations of the same stimulus (Figure 3A). Given that downstream decision-making neuronal ensembles likely pool information from many visually tuned neurons, we asked whether an optimal population decoder could reliably extract information about the visual stimulus across trials. We proceeded to build various logistic regression classifiers to decode which visual stimulus was presented from the brain-wide neuronal activity pattern during the stimulus presentation period of each trial. We found that despite the observed neuronal variability, the stimuli could be reliably decoded from the visually-tuned ROIs identified within the brain-wide population activity (Figure 3B) at the single trial level with an accuracy of 94 ± 2% (mean ± 95% confidence interval). This robust decodability suggests that the observed neuronal variability does not severely limit the information encoding capacity at the population level.

**Figure 3:**
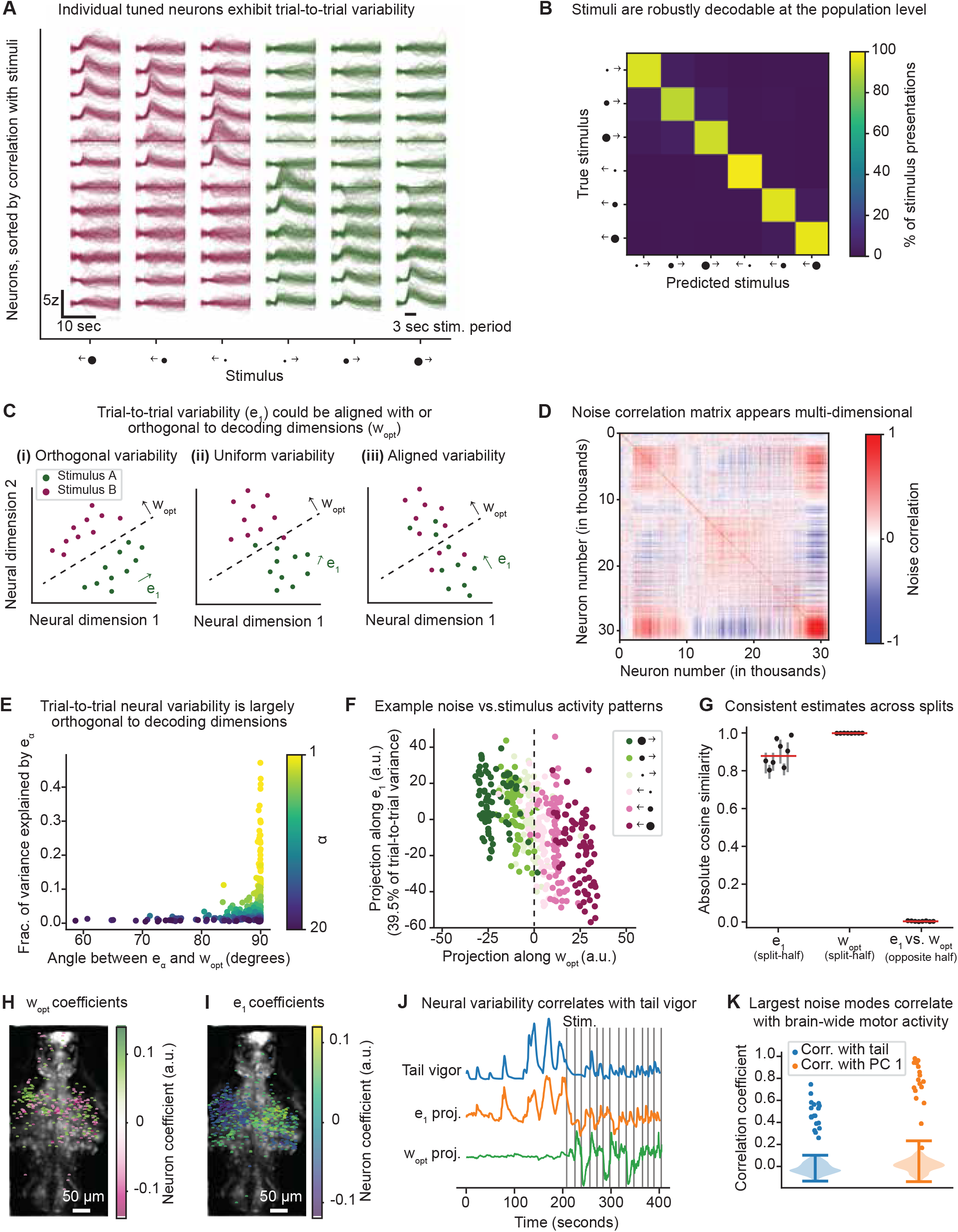
Trial-to-trial variability in visually-evoked neurons is largely orthogonal to visual decoding dimensions. **A.** Individual visually tuned neuronal ROIs exhibit trial-to-trial variability in their responses to stimuli. For each of the six visual stimuli (three object sizes on both the left and right visual fields), the two neurons exhibiting the highest correlation with the stimulus kernel (see Methods for details) are shown for an example larva. Each column represents a neuron, and each line represents its response to a given stimulus during a single trial. To visualize the neuronal activity during a given trial while accounting for the delay and kinematics of the nuclear-localized GCaMP (NL-GCaMP) sensor, a duration of approximately 15 seconds is extracted around each trial and the stimulus (“3 sec stim. period”) is indicated. Each trial is offset-corrected by subtracting the neuron’s baseline activity level at stimulus onset. **B.** Visual stimuli are reliably decodable from brain-wide dynamics on the single trial level. The visual stimuli were decoded using a one-versus-rest, multiclass logistic regression classifier with lasso regularization (see Methods for details). For each larva, a confusion matrix is computed for the test trials during 6-fold cross-validation. Shown is the average confusion matrix across n=8 larvae which were shown six visual stimuli. **C.** Schematic of potential geometric relationships between sensory decoding and neural variability dimensions. In each plot, each dot represents the neural response during a single presentation of stimulus A or B. The decision boundary for an optimal classifier is denoted with a dashed line, and the optimal stimulus decoding direction is denoted by the vector 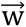_opt_. The direction representing the maximal trial-to-trial variance is denoted by 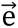_1_, and can be calculated by finding the first eigenvector of the noise covariance matrix (see Methods for details). These vectors can be: (i) orthogonal, such that neuronal variability does not limit the stimulus decoding; (ii) show little relationship, for example in the case of uniform variability; or (iii) aligned, such that variability likely limits the information encoding capacity along 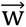_opt_. **D.** The trial-to-trial noise correlation matrix appears multi-dimensional. Shown is the average noise correlation matrix across all stimulus types presented. The ROIs are sorted using rastermap, which produces a one-dimensional embedding of the neurons, such that neurons which show similar correlation profiles are placed near to one another. A number of neuronal populations exhibiting correlations across trials are apparent from the clusters of high correlations near the diagonal. **E.** Trial-to-trial variability in the visually-evoked neurons is largely orthogonal to visual decoding dimensions. The fraction of trial-to-trial variance explained by each noise mode 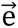_α_ is plotted against the angle between 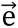_α_ and the optimal stimulus decoding direction 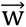_opt_. Shown are the 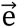_α_ for n=8 larvae, colored by their rank order α based on the fraction of variance explained. The largest noise modes were approximately orthogonal (∼90°) to the stimulus decoding direction, whereas only a few of the smallest noise modes exhibited angles less than 90°. **F.** Example projections of single trial neural activity along the stimulus decoding and noise dimensions. Each dot represents the average neural activity within a single trial projected along 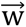_opt_ and 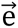_1_ for an example larva. Each of the six visual stimuli, three object sizes presented on either the right (green) or left (pink) visual field, are robustly encoded along 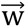_opt_ across trials; however, in all stimuli there is strong orthogonal variability along 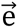_1_, the largest noise mode representing 39.5% of the trial-to-trial variance. **G.** 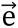_1_ and 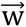_opt_ are reproducible and remain orthogonal across non-overlapping trial subsets. For each larva, trials within every stimulus condition were repeatedly divided into two non-overlapping 50:50 halves, and 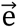_1_ and 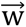_opt_ were estimated separately from each half. Absolute cosine similarity was calculated between the two 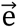_1_ estimates (“split-half”), between the two 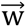_opt_ estimates (“split-half”), and between 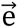_1_ and 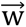_opt_ estimated from opposite halves (“opposite-half”). Each dot shows the median across 100 splits for one larva, and whiskers show the within-larva interquartile range. Red bars indicate the median across n=8 larvae. Values near 1 indicate alignment, whereas values near 0 indicate orthogonality. **H.** Example ROI coefficients for 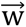_opt_. Shown are the 366 ROIs with the largest weights over a maximum intensity projection of the recorded volume. The visually-evoked ROIs which encode stimulus information are concentrated within the optic tectum. **I.** Example ROI coefficients for 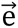_1_. Shown are the 828 ROIs with the largest weights over a maximum intensity projection of the recorded volume. The visually-evoked ROIs contributing to the largest noise mode are highly overlapping with the ROIs contributing to 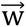_opt_ in panel H. **J.** The neural noise modes are highly correlated with tail movements. Shown are both the GCaMP kernel-convolved tail vigor and the neuronal projection onto the first noise mode 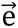_1_ for a representative larva. Over the full two-hour recording, the tail vigor and noise mode projection exhibit a significant correlation of r=0.58, p<10^-6^. For comparison, 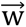_opt_ and the start time of the visual stimuli (black vertical lines) are shown. **K.** The largest neural noise modes reflect brain-wide motor encoding. Shown are the correlations between the first noise mode 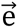_1_ and the tail vigor (blue) or the first principal component (PC 1, orange) of brain-wide data (see Figure S3C). Dots show data from individual larvae, whereas the violin plots below show the null distribution for temporally shuffled data, in which the tail vigor or PC 1 are circularly permuted.

Given the wealth of literature debating whether trial-to-trial noise correlations could interfere with stimulus encoding (Moreno-Bote et al., 2014; Kanitscheider et al., 2015; Zylberberg et al., 2017; Bartolo et al., 2020; Rumyantsev et al., 2020; Valente et al., 2021; Kafashan et al., 2021) and potentially drive behavioral variability, we next asked how the observed neuronal variability was structured to prevent degradation of the visual encoding. We began by considering the dimensions that maximally explain trial-to-trial neuronal variability within the high-dimensional space of neuronal population dynamics, which are sometimes referred to as “noise modes” (Rumyantsev et al., 2020). Historically, the term “noise” is used to describe such trial-to-trial variability that is not related to the external stimulus; however, such noise could represent significant internal neuronal dynamics and not just measurement shot noise, for example. If the primary noise modes (represented by neural activity along the vectors 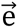_α_) are orthogonal to the optimal stimulus decoding direction (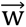_opt_), then trial-to-trial variability will not hinder the visual stimulus decoding (Figure 3C(i)). In this case, the neuronal population may carry a multi-dimensional set of at least two variables and the visual stimulus encoding would be preserved. Trial-to-trial variability could also not be limited to a particular set of dimensions, for example if it represented noise that is uniformly distributed in the neural state space (Figure 3C(ii)), in which case pooling or averaging across sufficient neurons may still recover adequate stimulus information. Finally, trial-to-trial noise modes may be aligned with the stimulus decoding dimension (Figure 3C(iii)), which would maximally degrade the visual stimulus decoding.

To assess which of these scenarios applied to the geometry of trial-to-trial neural variability in our data, we first utilized partial least squares (PLS) regression to identify a low-dimensional subspace of the brain-wide neural activity that optimally preserved information about the visual stimulus (Figure S3A, see Methods for details), which we refer to as the visually-evoked neuronal activity patterns. Importantly, this approach ensures that we are identifying trial-to-trial neural variation that is contained within the visually-evoked neuronal ROIs, as opposed to variability that is completely unrelated to visual encoding. Within these visually-evoked ROIs, PLS regression also identifies the optimal stimulus decoding direction 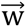_opt_ (Figure S3B). Additionally, trial-to-trial variability of the visually-evoked ROIs is summarized by their noise covariance matrix, which describes how the activity of neurons covaries across trials (Moreno-Bote et al., 2014; Kohn et al., 2015). The average noise correlation matrix across all stimuli (see Methods for details) appeared highly structured and multi-dimensional (Figure 3D), indicating the presence of multiple neuronal ensembles whose activity is strongly correlated across trials. We analyzed the structure of this trial-to-trial noise by finding the eigenvectors 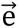_α_ of the average neural noise covariance matrix across all stimuli, following (Rumyantsev et al., 2020). As such, these noise modes 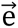_α_ represented the dimensions of neuronal activity within the visually-evoked ROIs that maximally covaried across trials. We found that the noise modes were strongly structured since the trial-to-trial variance was concentrated in relatively few dimensions. The single largest noise mode captured up to 50% of the variance across larvae (Figure 3E, y-axis), indicating that such activity is likely not merely “noise” but potentially physiologically relevant. Thus, the observed neuronal variability was not independent across ROIs, but strongly correlated. To assess the impact these noise modes could have on the stimulus decoding, we calculated the angles between the noise modes 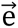_α_ and 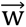_opt_. We discovered that the noise modes containing the majority of the variance were orthogonal to 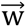_opt_, while only the smallest noise modes contained any overlap with the stimulus decoding direction 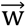_opt_ (Figure 3E).

This finding demonstrates that the structure underlying trial-to-trial neuronal variability in the zebrafish visual system lies in a regime closely resembling the orthogonal variability as illustrated in Figure 3C(i), as also previously observed in a subregion of mouse primary visual cortex (Rumyantsev et al., 2020). Thus, while even a small degree of observed overlap between sensory and noise modes can in principle limit the information encoding capacity of the visually-evoked neurons (Moreno-Bote et al., 2014), this orthogonal structure minimizes this limitation and is consistent with our previous finding that the stimuli presented in this study are highly decodable from the neuronal population dynamics (Figure 3B). In addition, while the angle between the sensory and noise modes arises from the pattern of coefficients across neuronal ROIs, we can visualize activity along these modes by projecting average neural activity within each trial onto the stimulus decoding direction 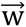_opt_ and the noise modes (Figure 3F). This demonstrates that the majority of the neural variability across trials (given by the projection onto 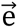_1_) is largely independent from the sensory activity (along 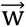_opt_).

Because these directions are defined in a multidimensional space spanned by the *d* PLS components, an approximately orthogonal angle alone is not sufficient to establish that their relationship is meaningful: random vectors become increasingly likely to appear orthogonal as dimensionality increases. We therefore used a repeated split-half analysis (see Methods for details) to determine whether 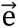_1_ and 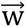_opt_ were individually reproducible and remained orthogonal when estimated from non-overlapping trials (Figure 3G). Within each recording, trials were stratified by stimulus condition and repeatedly divided into two non-overlapping halves. A common PLS basis was used so that directions estimated from the two halves were expressed in the same coordinate system; within this basis, 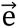_1_ and 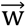_opt_ were estimated separately from each half. The median of the per-larva split-half absolute cosine similarities was approximately 0.88 for 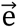_1_ and approximately 1.0 for 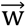_opt_. In contrast, the median cross-held-out absolute cosine similarity between 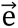_1_ and 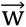_opt_ was approximately 0.003. Thus, both directions were reproducible across trial subsets, while their relationship remained nearly orthogonal.

Having established that this geometry was reproducible across trial subsets, we next asked whether the observed neuronal variability could be explained predominantly by eye movements, such as saccades or convergence. To do so, we tracked the angle of each eye. We utilized DeepLabCut, a deep learning tool for animal pose estimation (Mathis et al., 2018), to track keypoints on the eye which are visible in the raw fLFM images, including the retina and pigmentation (Figure S3D(i)). This approach enabled identification of various eye movements, such as convergence and the optokinetic reflex (Figure S3D(ii-iii)). Next, we extracted various eye states, including those based on position (more leftward vs. rightward angles) and speed (high angular velocity vs. low or no motion). Figure S3E(i) provides example stimulus response profiles across trials of the same visual stimulus in each of these eye states, similar to a single column of traces in Figure 3A broken out into more detail. These data demonstrate neuronal variability is still observed when controlling for eye state. We next asked more directly whether the dominant neural noise modes vary across eye states, which would suggest that the geometry of neuronal variability is dominated by eye movements or states. To do so, the dominant noise modes were estimated in each of the individual eye conditions. The similarity between noise modes estimated from different eye conditions (Figure S3E(ii), red distribution, right) largely overlapped the distribution obtained from resampled trial subsets pooled across all eye conditions (black distribution, left), although the eye-conditioned distribution was somewhat broader. Thus, although eye movements may contribute to the observed neuronal variability, they do not appear to dominate the geometry of the leading noise mode.

Although 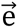_1_ and 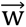_opt_ define distinct and nearly orthogonal directions in neuronal state space, this does not establish whether they arise from anatomically distinct neuronal populations. We therefore mapped these neural coefficient vectors defining the stimulus decoding direction and the noise modes onto their anatomical locations of the larvae’s brain. As expected, the visually-evoked ROIs were highly concentrated within the midbrain and specifically the optic tectum, such that both the stimulus decoding direction (Figure 3H) and noise modes (Figure 3I) highly overlapped. This confirms that the noise modes indeed represent fluctuations within the visual circuitry, as opposed to irrelevant activity from other brain areas. The significance of each ROI’s contribution to 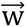_opt_ and 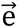_α_ was assessed by comparing its coefficient to a distribution of coefficients derived from trial-shuffled data (see Methods for details). Overall, 30 ± 6% of visually-evoked ROIs that significantly contributed to the largest noise mode 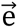_1_ also exhibited significant contributions to the stimulus decoding direction 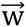_opt_ (mean ± 95% CI across n=10 larvae). Therefore, while many visually-evoked ROIs have response components along both 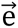_α_ and 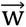_opt_, the neuronal population dynamics are highly structured such that activity patterns encoding external visual stimuli are reliable and largely independent from any trial-to-trial noise modes. Given that our data do not establish that every ROI corresponds to a single neuron, the observed orthogonality demonstrates separation between population-level coactivation patterns at the spatial resolution provided by fLFM, without excluding the possibility of functional heterogeneity within some ROIs.

Given that these noise modes represented highly-structured neuronal variability across trials, we importantly asked whether they were at all related to the larvae’s behavior. First, we asked whether any noise modes were correlated with the vigor of the larvae’s tail movements, defined as the absolute tail curvature time series convolved with the GCaMP response kernel (see Methods for details). We found that the largest noise modes were often correlated with the larvae’s instantaneous tail vigor (Figure 3J), as well as other kinematic variables (Figure S4A), for example tail curvature (which includes a left versus right directedness). While we identified the noise modes within the visually-evoked neuronal population, these noise modes were also highly correlated with the largest principal component (PC) of the brain-wide dynamics (Figure 3K). This indicates that both the observed trial-to-trial neuronal variability in visual regions and elsewhere across the brain are dominated by behavior-related activity (Figure S4B). In fact, 35 ± 15% of neuronal ROIs with significant coefficients in the largest visual noise mode 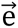_1_ also significantly contributed to the first brain-wide PC (mean ± 95% CI across n=10 larvae, see Methods for details). While the lower variance noise modes did not exhibit clear motor-related activity, we suspected they may be involved in additional internal computations that may modulate decision making across trials. Taken together, these results show that visual stimuli are faithfully represented on the population level, and that the observed behavioral variability is not driven by changes in the fidelity of sensory encoding across trials. Instead, behavioral variability may be explained by additional, orthogonal dimensions of neuronal activity either within the visually-evoked population or elsewhere in the brain that are stimulus-independent but predictive of the larvae’s behavior.

### Pre-motor neuronal populations predictive of single-trial behavior

We hypothesized that there exists a pre-motor neuronal population that would be predictive of the larvae’s behavioral response in any given trial *before* movement initiation. As such, we aimed to identify any neural locus predictive of the larvae’s turn direction at various timepoints through each visual stimulus presentation. Thus, we looked beyond only the visually-evoked population and returned to the brain-wide activity patterns. In order to build predictive models from the neuronal dynamics despite varying reaction time across trials, we applied time-warping to the neuronal timeseries within each trial such that the stimulus onset and movement initiation timepoints were aligned across trials (Figure 4A, see Methods for details). We selected trials in which the larvae had a reaction time of at least one second from the stimulus onset (providing sufficient pre-motor neuronal data) and made its first tail movement with a defined minimum tail curvature to either the left or right, so as to separate clear left and right turns from simple forward locomotion. Further, we utilized trials from all visual object sizes to collect enough trials to reliably train models and thus identify populations predictive of decisions regardless of incoming sensory information. We found that there appeared to be neuronal ROIs with differing time-warped activity patterns during left and right turn trials (Figure 4A, bottom). Additionally, we extracted trials in which the larva did not respond following the stimulus presentation, which were time-warped according to a randomly selected reaction time from the distribution in responsive trials. As such, each trial could be categorized according to the stimulus presented, the larva’s responsiveness (response/no response), and its turn direction (left/right) within responsive trials.

**Figure 4:**
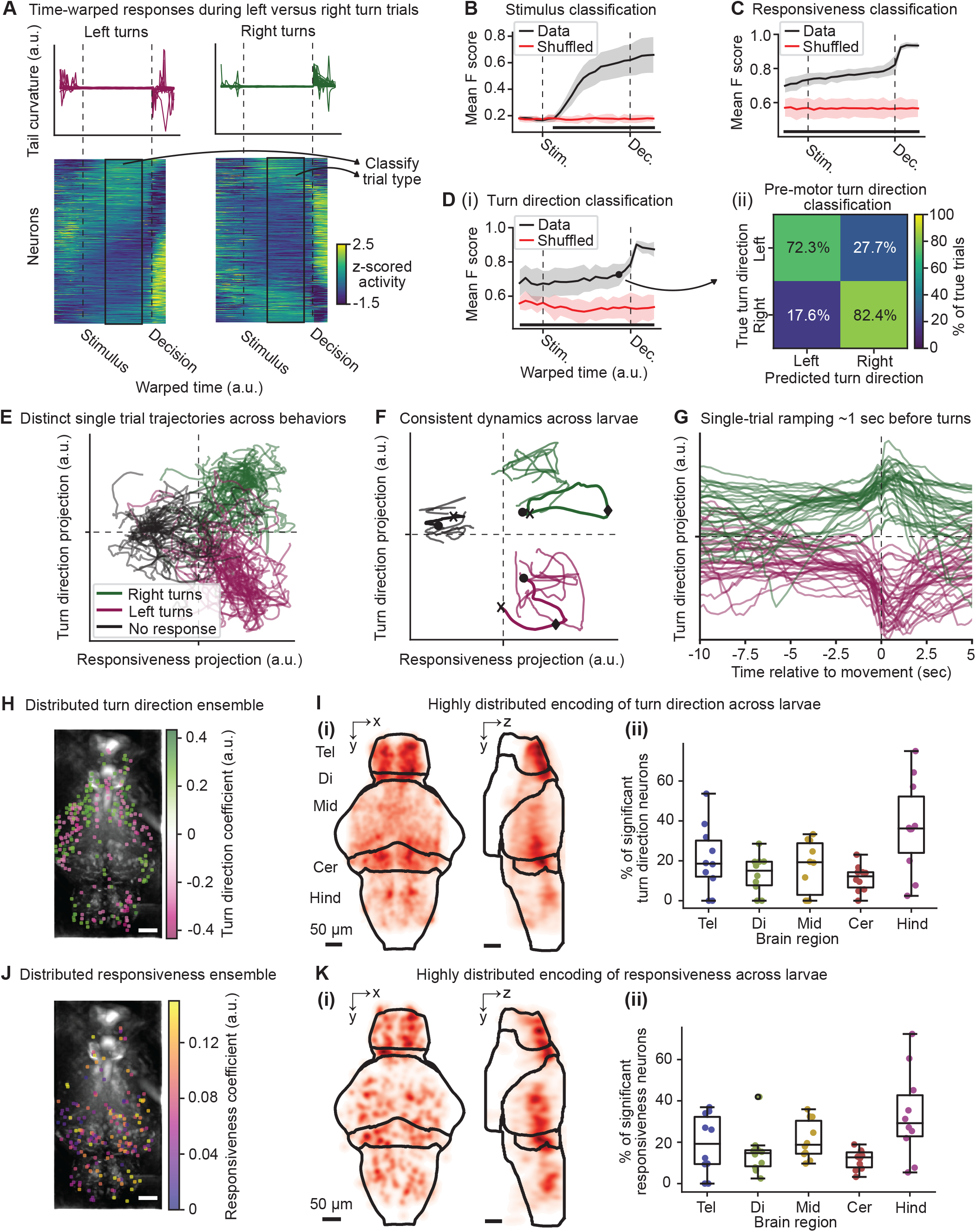
Pre-motor neuronal populations predictive of single-trial behavior. **A.** Trials are time-warped to align stimulus and decision onsets before classifying the turn direction. Top: Time-warped tail curvatures for trials in which the fish performed leftward or rightward turns, on the left and right, respectively. Bottom: Trial-averaged and time-warped neuronal timeseries for 15,286 neuronal ROIs during left and right turns. The neurons are sorted using the rastermap algorithm. A time window is swept across the stimulus and decision timepoints to train binary classification models to predict the turn direction from the associated neuronal dynamics. **B.** Stimulus classification accuracy peaks after the onset of visual stimulation. The mean F score across n=7 larvae is used to assess the performance of 6-way multiclass classification of the presented visual stimulus as a function of warped time surrounding the stimulus onset (Stim.) and decision timepoint (Dec.). Shown is the mean ± 95% confidence interval of the F score for the best time window ending at the given timepoint (Data), compared to shuffled data in which the class labels are randomized. The black bar at the bottom indicates timepoints where the data have a significantly higher F score than the shuffled data (p<0.05, paired t-test). **C.** Binary classification of responsiveness, whether or not the fish responds in a given trial, is significant throughout all time periods but accuracy peaks near movement initiation. As in panel B, except for binary classification of responsiveness. Nonresponsive trials are time-warped by randomly selecting a reaction time from the response trials and applying the same transformation. **D.** (i) Turn direction classification accuracy is significantly higher than shuffled data across the entire time-warped interval, but peaks near movement initiation. As in panel B, except for binary classification of turn direction. (ii) Single trial classification of turn direction across larvae. The mean confusion matrix across n=7 larvae shows an accuracy of 77 ± 4% (mean ± 95% confidence interval). **E.** Single trial trajectories are separated based on responsiveness and turn direction. Shown are neural activity trajectories during single trials in an example larva projected onto the brain-wide neural dimensions that optimally separated turn direction and responsiveness. **F.** Consistent trial-averaged trajectories across larvae. As in panel E, except for the trial-averaged responses for n=6 example larvae. For the one bold animal, timepoints across the trial are indicated by a circle for trial start, diamond for the decision timepoint, and an X for the trial end. **G.** Real-time single-trial dynamics in an example larva. Along the turn direction neural projection, left and right trials are separated for many seconds before the decision timepoint, which is longer than the three second length of visual presentations. Activity along this dimension shows consistent ramping across trials approximately one second before movement. **H.** Example turn direction neuronal ensemble. Shown are the coefficients for all neurons which showed significantly higher (one-tailed t-test, p<0.05) absolute coefficients in the real models compared to shuffled data in which the turn direction labels are randomly permuted. Scale bar: 50 μm. **I.** Highly distributed encoding of turn direction across larvae. The significant turn direction neurons located within five major brain regions are shown for n=10 larvae. (i) The 3D distribution of neurons across larvae aligned to the Z-Brain atlas. (ii) Quantification of the percentage of turn direction neurons located within each brain region shown in (i). There is no significant difference between the percentage of neurons across brain regions (p>0.05, paired t-test). **J.** Example responsiveness neuronal ensemble. As in panel H, except for responsiveness. Shown are the coefficients for all neurons which showed significantly higher (one-tailed t-test, p<0.05) absolute coefficients in the real models compared to shuffled data in which the responsiveness labels are randomly permuted. Scale bar: 50 μm. **K.** Highly distributed encoding of responsiveness across larvae. As in panel I, except for responsiveness. There is no significant difference between the percentage of neurons across brain regions (p>0.05, paired t-test).

We first asked whether the larvae’s responsiveness and turn direction could be accurately classified from the brain-wide neuronal dynamics within single trials, as well as how this predictability behaved over the pre-motor period. To identify the timepoints throughout the trial within which these could be accurately classified, we varied the time window of neuronal dynamics used to build a binary logistic regression classification model (Figure S5A). Neuronal activity across all ROIs was integrated from the window start to the window end separately for each trial and then a model was trained on 80% of the trials. To assess the prediction accuracy at various timepoints throughout the trial, we utilized the mean F score across larvae on held-out trials, which measures the harmonic mean of the precision and recall. As expected, the multiclass stimulus prediction accuracy peaked shortly after the stimulus onset and remained high throughout the pre-motor and decision periods (Figure 4B). Performing stimulus classification again using only neuronal ROIs from individual brain regions, we found that only populations in the midbrain and cerebellum exhibited strong stimulus decoding (Figure S5B). For both the responsiveness (Figure 4C) and the turn direction (Figure 4D(i)), the mean F score was consistent across the pre-stimulus and stimulus periods, before quickly ramping near the movement initiation timepoint. Interestingly, the classification performance was significantly higher than shuffled data at all timepoints (Figure 4C-D), including before the start of the stimulus presentation, with an average accuracy of 77.4 ± 4.4% (mean ± 95% CI across n=7 larvae) during the pre-motor period (Figure 4D(i)). These results suggest there are two dominant timescales which contribute to the larva’s behavioral response: a longer-timescale and relatively weak (but significant) activity pattern that does not appear aligned to stimulus or behavioral events, which we term the pre-stimulus turn bias; and a fast-timescale dramatic increase in predictability very near the decision, which we term the movement initiation dynamics.

Given that we observed the largest degree of behavioral variability in the responses to small-to-intermediate size stimuli (Figure 2G), we hypothesized that the pre-stimulus turn bias would be most influential (and thus most predictive of behavior) during such stimulus presentations, and less so during presentations of larger stimuli. Indeed, while the mean F scores for turn direction prediction were highly variable across larvae, on average the largest stimuli exhibited the lowest predictability from the pre-stimulus turn bias signal (Figure S5C, blue line). This likely reflects a more salient stimulus drive during large stimulus trials, which in the natural environment could reflect predators and a potential life-or-death decision for the larvae. However, significant predictability was observed across all stimulus sizes during the movement initiation dynamics (Figure S5C, orange line). We thus argue that the pre-stimulus turn bias is subsequently combined with incoming stimulus information during action selection. Ultimately, the larva’s decision is then signaled by the dramatic increase in predictability during the movement initiation dynamics.

These single-trial predictions could be visualized by projecting the brain-wide neuronal activity during each trial onto the neural dimensions that optimally classified turn direction and responsiveness. Consistent with our classification performance, we found that the trajectories during single trials (Figure 4E) were highly separated according to the ultimate behavioral response. On average and across larvae, these trajectories remained localized in distinct quadrants throughout the entire trial (Figure 4F), including before and during the stimulus presentation period. This suggests that the observed behavioral variability is due in part to the initial internal state of the neural dynamics at the time of stimulus presentation and decision. Indeed, by removing the time-warping and observing the dynamics during single trials in real time, left and right turn trials appear separated for many seconds prior to the movement (longer than the 3 second visual trials). Ultimately, the decision to turn during the movement initiation dynamics appeared driven by a strong ramping activity approximately one second before the turn (Figure 4G). Thus, the identified neuronal dimensions exhibited reliable decoding within single trials driven by an initial bias and ultimately resulting in the observed movement initiation dynamics.

Next, we asked which specific neuronal populations across the brain contributed to the accurate pre-motor classification of behaviors, and thus could be involved in driving the decision of when and how to act. We considered a neuronal ROI to be significantly contributing to the behavioral prediction if its distribution of absolute coefficients across 6-fold cross-validation was significantly higher (one-tailed t-test, p<0.05) when fit to real behavioral data as opposed to a null distribution created by randomly permuting the trial type labels. We found that both the ensemble of neurons which significantly contributed to the turn direction prediction (Figures 4H-I) and the responsiveness ensemble (Figures 4J-K) were highly distributed across the brain, with potentially dense localization in the telencephalon, cerebellum, and dorsal diencephalon (habenula). Both of these distinct neuronal ensembles showed no significant difference in their distribution across major brain regions: the telencephalon (Tel), diencephalon (Di), midbrain (Mid), cerebellum (Cer), and the remaining hindbrain (Hind). Further, while this population included visually-evoked ROIs within the optic tectum, we found that our previously identified noise modes 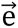_α_ were similarly predictive of single trial turn direction (Figure S5D), whereas the optimal stimulus decoding direction 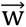_opt_ (Figure 3H) was not able to predict single-trial turn direction (Figure S5E).

Finally, given that our turn direction and responsiveness classification required using lasso regularization to induce sparsity and achieve strong prediction performance, we aimed to identify whether a larger selection of neurons may contribute to or reflect these activity patterns by quantifying the correlation between each ROI and the projection of the brain-wide neural activity along the optimal turn direction or responsive prediction directions. An ROI was considered significantly correlated with either projection if its correlation was significantly higher than that of shuffled data obtained by circularly permuting the ROI’s timeseries (see Methods for details). These distributions (Figures S5F-G) strongly resembled those in Figures 4H-K and supported the brain-wide nature of these representations. Further, the predictability of turn direction or responsiveness was similar across these brain regions (Figures S5H-I). Thus, our data suggest that when stimuli are ambiguous, single-trial action selection is largely explained by a widely-distributed circuit containing subpopulations encoding internal time-varying biases related to both the larva’s responsiveness and turn direction, yet distinct from the sensory encoding circuitry.

What could be the origin of these trial-to-trial biases in the larvae’s behavior? Given that our results demonstrate that it is possible to predict a behavioral bias before the visual stimulus is shown, we hypothesized these results could reflect the influence of the zebrafish neuronal circuitry for spontaneous turns, which includes the hindbrain oscillator (or anterior rhombencephalic turning region, ARTR) known to bias sequential turns towards a given direction (Dunn et al. 2016b). While our pre-motor neuronal population is not exclusively localized to the ARTR, we did identify many neuronal ROIs in the cerebellum and hindbrain (Figures 4J-K) whose locations were consistent with these known populations. As such, we asked whether spontaneous turns could be predicted from this same turn direction neuronal ensemble (Figure 5A). To address this question, we selected spontaneous turns that occurred either during inter-trial intervals or within a 2-minute period at the beginning of each recording without any stimulus presentations, utilizing the same criteria to define left and right turns as previously (see Methods for details). We then asked whether the spontaneous turn direction during periods without sensory stimulation could be predicted from the same turn direction neuronal ensemble and coefficients previously identified in Figure 4H-I. We found a significant correlation between the pre-motor turn direction predictions and the observed spontaneous tail curvature across larvae (Figure 5B, r=0.41, p<0.05), suggesting this same ensemble is activated prior to both spontaneous and visually-evoked turns. This ensemble had a pre-motor turn direction classification accuracy of 70.2 ± 6.0% (mean ± 95% CI across n=5 larvae, Figure 5C). Thus, these results demonstrate that a portion of the observed behavioral variability is overlapping with the larva’s motor circuitry responsible for generation of spontaneous behavior.

**Figure 5:**
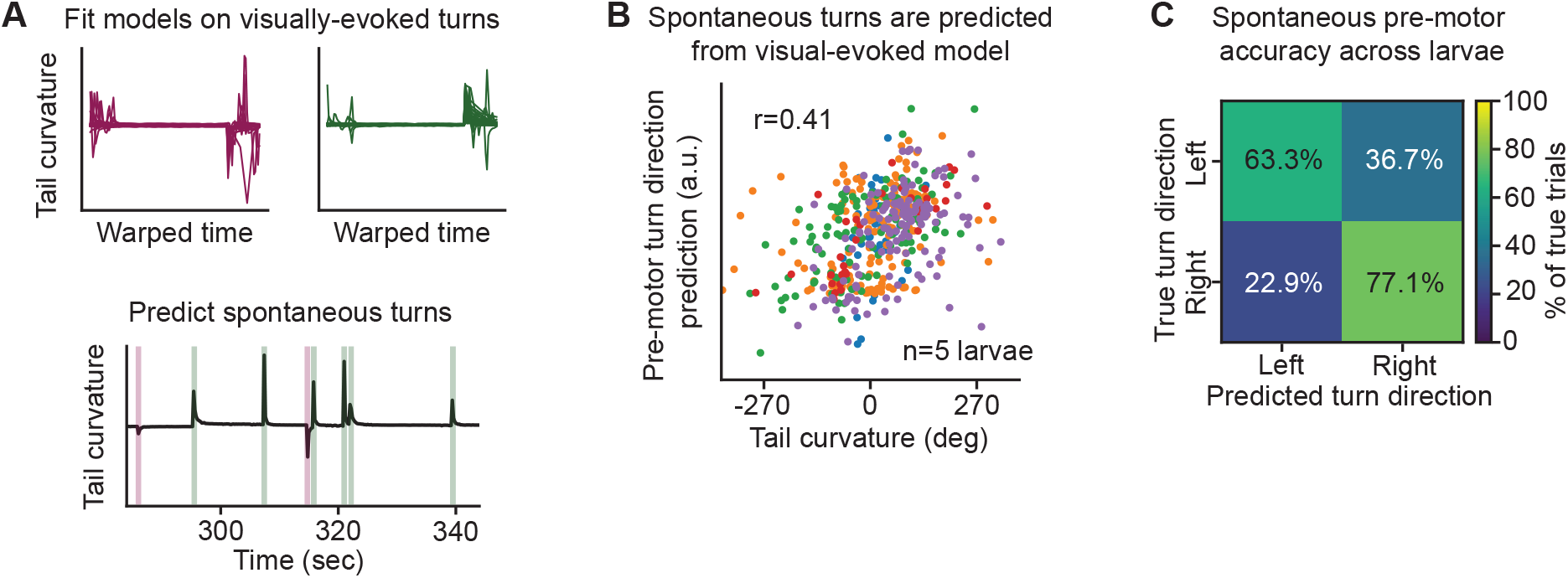
Spontaneous turns are predictable from the same pre-motor neuronal population. **A.** Schematic of the approach to predict spontaneous turns. Models are fit to predict left or right turn direction during visual trials (top), as in Figure 4. They are then tested on pre-motor periods one second before the spontaneous turn. **B.** Spontaneous turns are predicted from the visual-evoked model. Shown is the relationship between the spontaneous tail curvature and the predicted pre-motor turn direction using the visual-evoked model. Each dot is a single spontaneous turn and each color represents a different larva. They exhibit a significant correlation of r=0.41, p<0.05. **C.** Spontaneous turn classification accuracy. The mean cross-validated confusion matrix for spontaneous turn classification over n=5 larvae. Spontaneous turns are predicted with an accuracy of 70.2 ± 6.0% (mean ± 95% CI across n=5 larvae).

Our data highlight that the neural mechanisms involved in single-trial decision making are reflected in a highly distributed ensemble of neurons across the larval zebrafish brain. Further, behavioral variability, particularly in the larvae’s responses to small-to-intermediate size stimuli, can be partially explained by an internal, task-independent turn bias that is unrelated to the visual stimulation, while their stimulus-driven responses to larger predator-like stimuli exhibited a comparatively lower level of behavioral variability. This suggests the importance of considering how internal biases and circuits governing spontaneous movements interact with the well-studied sensorimotor circuits to govern how the brain generates behavior in a naturalistic context.

## DISCUSSION

In this study, we designed and built an optimized, high-resolution Fourier light field microscope (fLFM) to perform high-speed volumetric imaging across the larval zebrafish brain and investigate trial-to-trial variability in large, anatomically distributed neuronal populations. To do so, we studied visually-evoked turns within two ethologically relevant behaviors with opposing valance, prey capture and escape response, each of which is on average driven by distinct pre-motor neuronal ensembles (Bollmann, 2019) dependent on the size of the given sensory stimulus. Consistent with previous results (Barker and Baier, 2015), we found that the larvae’s behavioral responses were highly variable across presentations of the same stimulus, with behavioral variability peaking in responses to stimuli of an intermediate size between those which optimally drive attraction or avoidance behaviors. Given that we lack a mechanistic understanding of which specific neuronal circuits drive the observed variance across trials, we utilized fLFM to screen brain-wide population dynamics at the level of individually segmented neuronal ROIs and identify neuronal ensembles which contribute to behavioral variability at the single-trial level.

We first asked whether the observed behavioral variability could be explained by noise or trial-to-trial deviations in encoding of the visual stimuli. We found that despite the variable responses of individual neuronal ROIs to visual stimuli across trials, the population response faithfully and reliably encoded the visual stimulus in each trial. Further, we discovered that the visually-evoked neuronal activity patterns were nearly orthogonal to those dimensions that maximally covaried across stimulus repetitions, i.e. the noise modes, indicating that the behavioral variability was not explained by unreliable representation of the sensory inputs. Instead, we found that at least a third of the visually-evoked ROIs contributed to noise modes that were correlated with various motor outputs. We ultimately identified two brain-wide neuronal populations which could predict the larvae’s turn direction and responsiveness at the single-trial level. Surprisingly, a pre-stimulus bias component of this neuronal population’s activity could predict the larvae’s trial-by-trial turn direction even before the onset of the stimulus presentation with an average accuracy of 77%. After stimulus onset, a sharp ramping of the activity of this neuronal population approximately one second before movement initiation allowed for an increased turn direction prediction accuracy of 90%. Taken together, our data show that the larva’s trial-by-trial decisions in response to any given stimulus are partially explained by pre-stimulus turn biases, and then ultimately driven by movement initiation dynamics which are broadcast within brain-wide neuronal ensembles.

In this context, the design of our behavioral paradigm has allowed us to gain insights into the nature of trial-to-trial neuronal and behavioral variability within and across two different ethologically relevant but highly divergent behaviors by tuning only a single parameter. This has allowed us to show that functionally different neuronal circuits have evolved to exhibit different levels of variability. Further, the differential interaction of such neuronal variability with the well-studied sensorimotor circuits may govern how the brain generates optimal behavior in different naturalistic contexts.

In addition, our study represents the first to our knowledge that identifies behavior-related noise correlations throughout an entire brain-wide sensorimotor decision-making circuit. The nearly orthogonal relationship between the stimulus decoding direction and trial-to-trial noise correlations is highly consistent with previous studies which found that noise correlations within single brain regions are organized so as to prevent additional information encoding channels within the same population from limiting sensory encoding capacity (Kafashan et al., 2021; Rumyantsev et al., 2020). However, even a small degree of overlap between noise and sensory dimensions may decrease the information encoding capacity of the population (Moreno-Bote et al., 2014). Why then might the brain favor a coding scheme in which populations mix information encoding with additional noise correlation structure?

Our data offer one possibility: that such additional correlations could encode internal or behavioral states that modulate sensorimotor transformations and behavioral output, enabling dynamic and flexible behavioral responses. Indeed, we found that a large fraction of the observed trial-to-trial neuronal variability was related to the larva’s behavior: including activity correlated with the instantaneous behavioral state of the animal as well as pre-stimulus turn biases that correlated with responsiveness and turn direction. We found that the largest trial-to-trial noise mode within the visually-evoked ROIs was a subset of a brain-wide population correlated with the vigor of the larva’s tail movements. This is conceptually similar to recent evidence in rodents that a large fraction of spontaneous neural activity reflects spontaneous and uninstructed movements (Musall et al., 2019; Stringer et al., 2019; Manley et al., 2024).

Taken together, our study represents the first brain-wide confirmation that behaviorally relevant information is highly mixed throughout neuronal populations involved in processing sensory and other information, potentially enabling flexible and context-dependent behavior. Our observation of time-varying neuronal ensembles encoding a turn direction bias is reminiscent of the larval zebrafish’s spontaneous swimming circuitry, which involves a central pattern generator (the ARTR) that biases the larva to swim in chains of turns in alternating directions (Dunn et al., 2016b); however, the neuronal populations we identified were not localized to the previously identified circuitry, and instead were distributed across the brain. While such spontaneous turn sequences are thought to underlie efficient exploration of local environments, our data could reflect a similar sensorimotor circuitry that injects variability as a learning mechanism to explore the space of possible sensorimotor consequences. Given the brain-wide distribution of the behavior-related ensembles within the variable visually-evoked behaviors studied here, we expect that they are not driven solely by a single central pattern generator as with ARTR and spontaneous turns, but instead involve the interaction of such pattern generators with the larva’s internal state and recent experience. It has been shown that such hindbrain oscillatory activity integrates recent visual information to enable efficient phototaxis (Wolf et al., 2017), suggesting these pattern generators could act in concert with additional contextual populations. For example, it has been shown that hunger shifts decisions from avoidance to approach due to neuromodulation from the hypothalamic and serotonergic systems (Filosa et al., 2016). Further, our observed dynamics could also reflect the foraging state of the larva, which is known to be encoded by an oscillatory, neuromodulatory network distributed across the brain (Marques et al., 2020) and is a likely candidate to modulate a larva’s response to prey-sized stimuli. We envision that utilization of the wide array of genetically labelled lines that have been associated with these sensorimotor and neuromodulatory circuits (Semmelhack et al., 2014; Barker and Baier, 2015; Marques et al., 2020; Okamoto et al., 2021) could disentangle the specific contributions of the sensory, pre-motor, and internal state circuitry within the brain-wide populations observed here. Of particular interest will be how the activity of these populations is organized over longer timescales and modulated by visual input or internal states related to hunger and foraging, which are critical to balance feeding behaviors and predator avoidance within the developing larvae.

## Supporting information

Reviewers Response Letter

## ACKNOWLEDGEMENTS

We thank Q. Lin and S. Otero Coronel for discussions and feedback on the manuscript, T. Nöbauer and Y. Zhang for discussions regarding optical designs, and the members of the Vaziri Lab for discussions regarding the experiments and data analysis. We also thank A. Kaczynska, S. Campbell, and J. Hudspeth for housing zebrafish, and the Rockefeller University High Performance Computing Cluster for compute access. This work was supported in part by the National Institute of Neurological Disorders and Stroke of the National Institutes of Health under award numbers 1RF1NS113251 and 1RF1NS110501 and the Kavli Foundation through the Kavli Neural Systems Institute.

## AUTHOR CONTRIBUTIONS

J.M. contributed to the project conceptualization, designed and built the imaging system, performed experiments, analyzed data, and wrote the manuscript. A.V. conceived, led and supervised the project, designed the imaging system and biological experiments, guided data analysis, and wrote the manuscript.

## FIGURE CAPTIONS

**Supplementary Figure S1:**
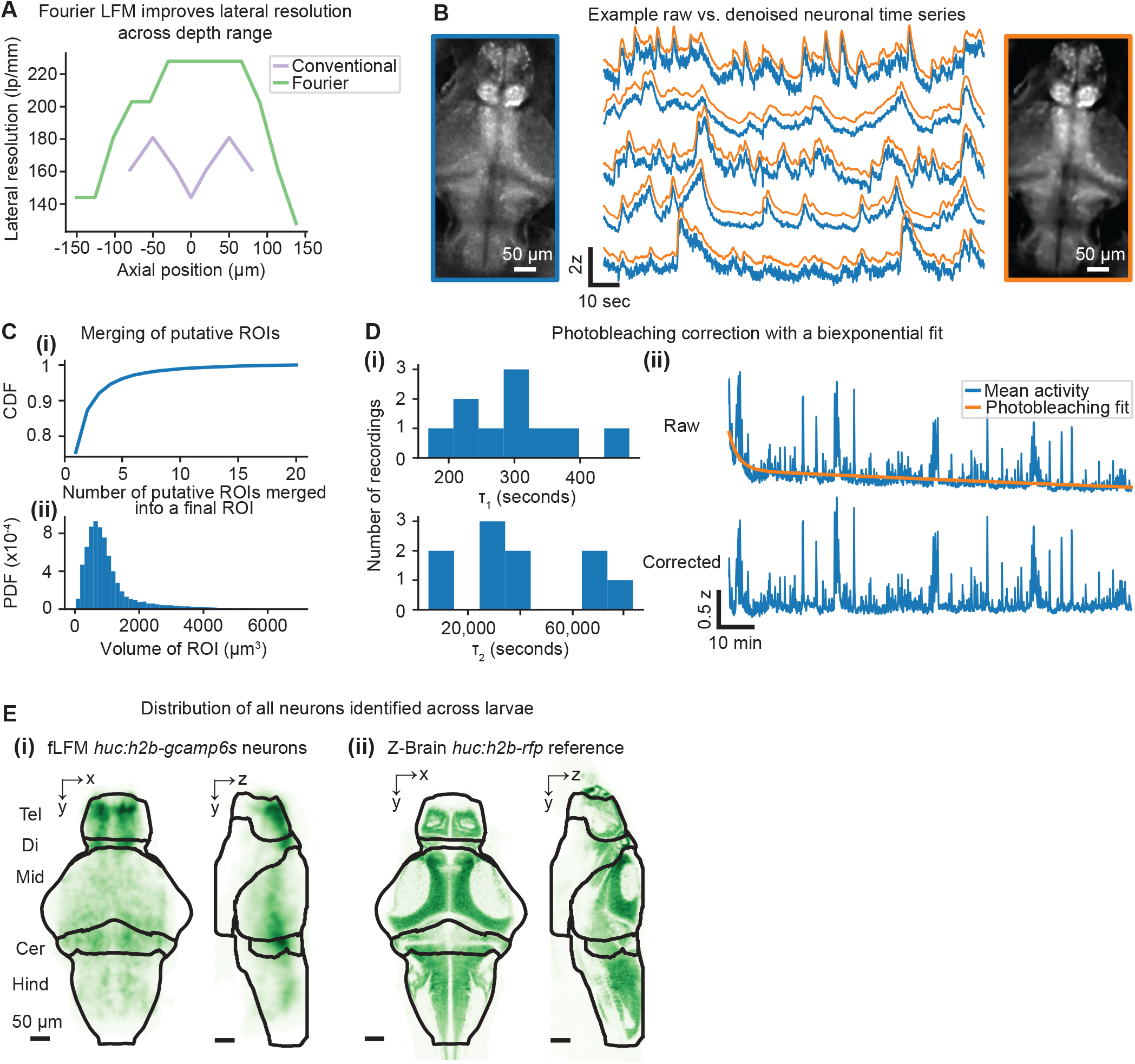
related to Figure 1, Fourier Light Field Microscopy (fLFM) provides a simple and cost-effective method for brain-wide imaging of larval zebrafish during behavior. **A.** Reconstructed lateral resolution of fLFM and cLFM across the axial imaging range, measured using a USAF resolution target translated relative to the native image plane. fLFM (green) exhibits higher lateral resolution and an expanded axial imaging range compared to cLFM (purple). **B.** Examples of raw and denoised neuronal time series. Left: an example reconstructed plane from a raw fLFM image. Blue circle indicates the ROI whose activity is plotted in the middle. Middle: Raw (blue) and denoised (orange) time series of average fluorescence intensity from neurons detected in the plane shown in the images. Right: the corresponding reconstructed plane as on the left, but after denoising the fLFM image with DeepInterpolation. **C.** Statistics of merging putative ROIs with high correlation and low spatial distance (see Methods for details) across n=10 larvae. (i) The cumulative distribution function (CDF) of the number of putative ROIs which were merged to create a final neuronal ROI. (ii) The probability density function (PDF) of the volume of the final neuronal ROIs. **D.** The effect of photobleaching on the fluorescence intensity was corrected with a biexponential fit. (i) The distribution of time constants resulting from biexponential fits to the mean neuronal signal for n=10 larvae. Two time constants are fit for each recording, one on the order of minutes (top) and the other tending to be on the order of many hours (bottom). (ii) Examples of raw (top) and corrected (bottom) mean neuronal signals from one recording. The biexponential fit to the data is shown on the top in orange. **E.** The distribution of neuronal ROIs across n=10 larvae resembles the labeling from the Z-Brain reference image. (i) The distribution of identified neurons (labeled with *huc:h2b-gcamp6s*) across all recordings shown aligned to the Z-Brain atlas. Black borders mark the boundaries of the telencephalon (Tel), diencephalon (Di), midbrain (Mid), cerebellum (Cer), and the remaining hindbrain (Hind). Shown are the xy and yz projections of the kernel density estimate of the center of mass of each ROI. (ii) One xy and yz slice from the *huc:h2b-rfp* reference image to which each fLFM recording was aligned. Scale bars represent 50 μm. The distribution of neuronal ROIs in the fLFM recordings is broadly similar to the reference image across major brain divisions; however, labeling and detected ROI density are reduced in deep ventral regions, including the hypothalamus. Improved pan-neuronal transgenic lines or more targeted labeling may be desired for these regions.

**Supplementary Figure S2:**
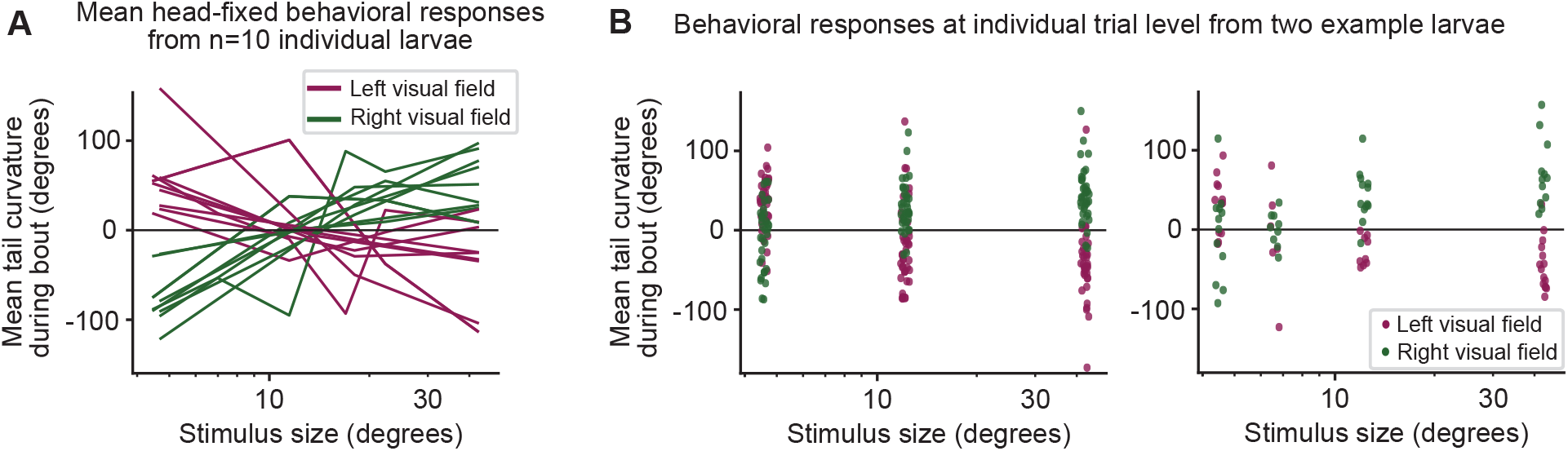
related to Figure 2, zebrafish exhibit highly variable motor responses to visual stimuli. **A.** Inter-individual variability in the motor responses demonstrated by the mean head-fixed behavioral response from n=10 larvae, as shown in Figure 2F but at the individual larva level. **B.** Intra-individual variability in the motor responses shown for two example larvae (left and right), similar to Figure 2G. Each point represents the mean tail curvature during a single trial.

**Supplementary Figure S3:**
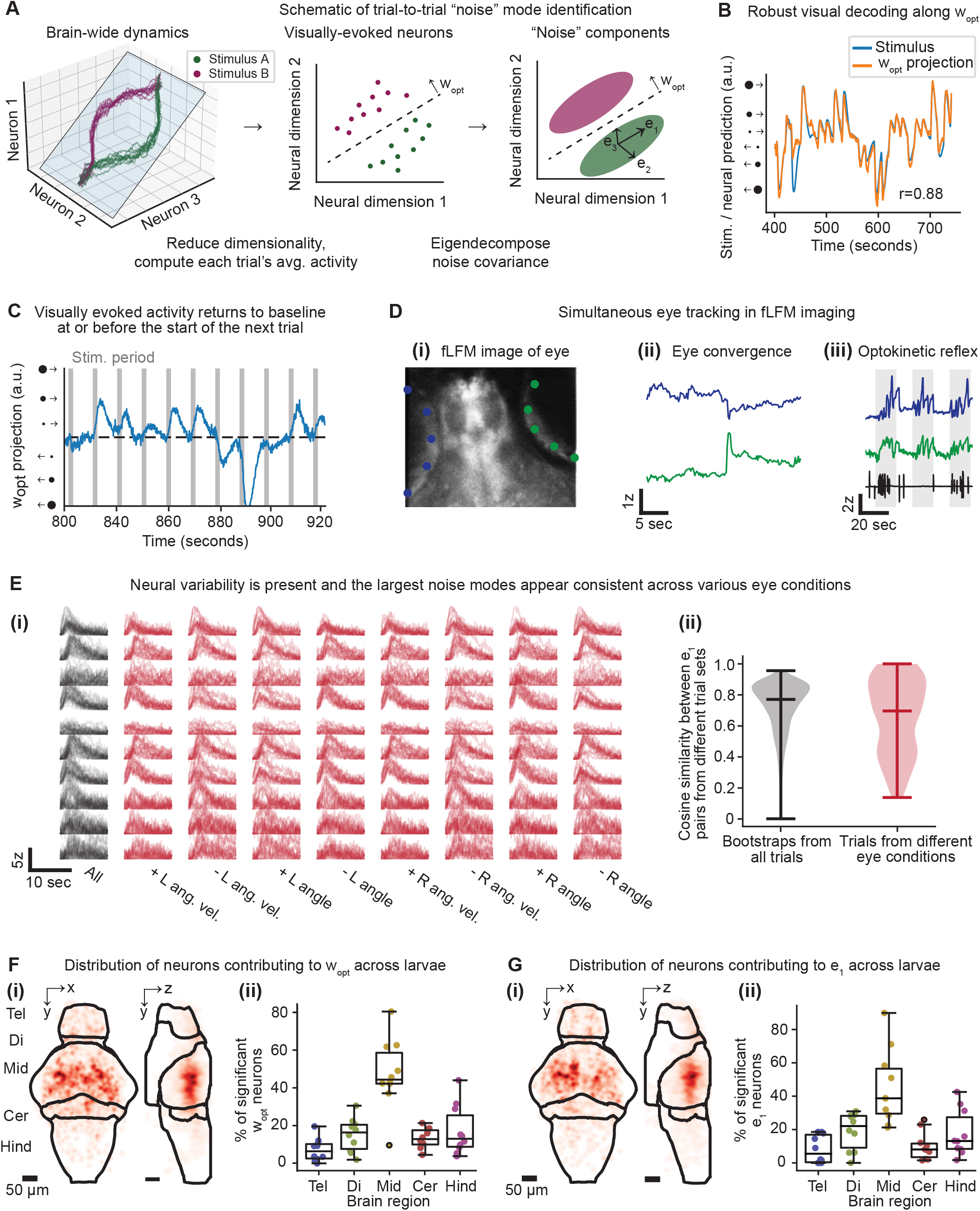
related to Figure 3, trial-to-trial variability in visually-evoked neurons is largely orthogonal to visual decoding dimensions. **A.** Schematic of trial-to-trial noise mode identification. First, dimensionality reduction is applied to the brain-wide dynamics (left) to identify the neuronal activity patterns that optimally preserve visual information using partial least squares regression. Within these visually-evoked neurons, each trial’s average activity during the visual presentation is computed (middle, each dot represents a single trial corresponding to the pink or green stimulus condition). Finally, the covariance across trials is decomposed to find the noise modes 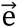_α_, which represent orthogonal directions within the visually-evoked neurons and are ranked in order of decreasing variance. **B.** 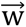_opt_ robustly encodes the visual stimulus. Shown are example time traces of the stimulus kernel (blue, where the size of the stimulus is encoded by the magnitude and which visual field it is presented in is encoded by the sign) and the neural projection onto 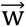_opt_ (orange), which exhibit a correlation of r=0.88, p<10^-6^. **C.** Visually-evoked activity returns to baseline at or before the start of the following stimulus presentation. The projection along the optimal stimulus decoding direction 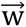_opt_, which demonstrates the dynamics of the visually-evoked population, is shown relative to the duration of the visual stimulus presentations (gray shaded bars). **D.** Tracking of various eye movements in the raw fLFM images. (i) An example raw fLFM image of an eye, with keypoints tracked in DeepLabCut shown for the left (blue) and right (green) eyes. (ii) An example eye convergence event. (iii) An example of the optokinetic reflex. Saccades, in which both eyes exhibit motion in the same direction, are shown as the larva tracks a moving grating stimulus (gray shaded regions). The tail curvature is shown below in black, demonstrating the optomotor reflex as well. **E.** Neural variability is consistent across various states of the eye and eye movements. (i) For one visual stimulus type, the activity of 10 neurons (rows) is shown during various subsets of trials (columns). While all of the trials for this stimulus are shown on the left in black, the remaining red traces exhibit various conditions for each eye (L or R): high (+) or low (-) angular velocity (ang. vel.), as well as larger, leftward looking eye angles (+ angle) or smaller, rightward looking eye angles (- angle). (ii) Comparison of the dominant noise mode estimates across trial subsets. A control distribution is shown on the left in black by performing bootstrapping with replacement to randomly select a subset of trials and then estimating 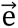_1_. On the right in red, noise modes are compared from the different eye conditions in (i). The noise mode estimates are compared using the absolute cosine similarity, where 1 indicates perfect alignment whereas 0 indicates orthogonality. **F-G.** Distribution of ROIs contributing to 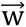_opt_ (F) or 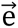_1_ (G) across n=10 larvae. (i) The 3D distribution of significantly contributing neurons, shown via xy and yz projections relative to the Z-Brain atlas. (ii) Quantification of the percent of contributing neurons contained within each of the brain regions shown in (i).

**Supplementary Figure S4:**
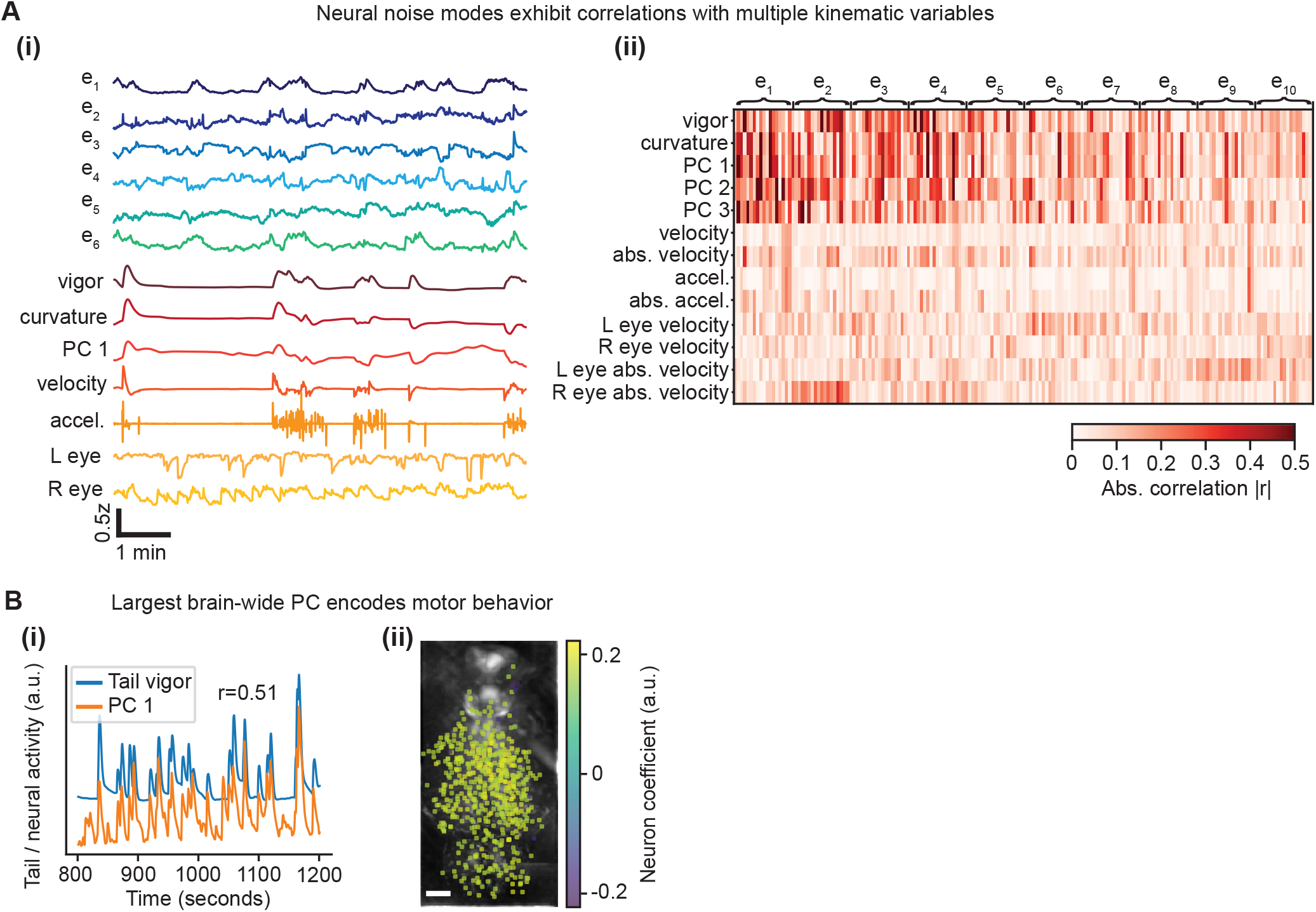
related to Figure 3, trial-to-trial variability in visually-evoked neurons is largely orthogonal to visual decoding dimensions. **A.** Multiple neural noise modes show correlations with motor outputs. (i) Example timeseries of six neural noise modes 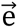_α_ and various kinematic variables: tail vigor, tail curvature, the first principal component (PC 1) of the tail angles, tail angular velocity, tail angular acceleration, left eye angular velocity, and right eye angular velocity. Each of the kinematic variables is convolved with the NL-GCaMP6s response kernel. (ii) Absolute correlation matrix between the neural noise mode and kinematic timeseries for n=10 larvae. Each row represents one motor variable. The columns are grouped by noise mode 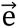_α_. Within a noise mode group, each individual column represents one larva. **B.** The largest brain-wide principal component (PC) encodes motor behavior. (i) Example time traces of the tail vigor (blue) and PC 1 (orange), which exhibit a correlation of r=0.51, p<10^-6^. (ii) The coefficients of the ROIs contributing most to the PC, which are distributed across the brain and are nearly all positively correlated with tail vigor.

**Supplementary Figure S5:**
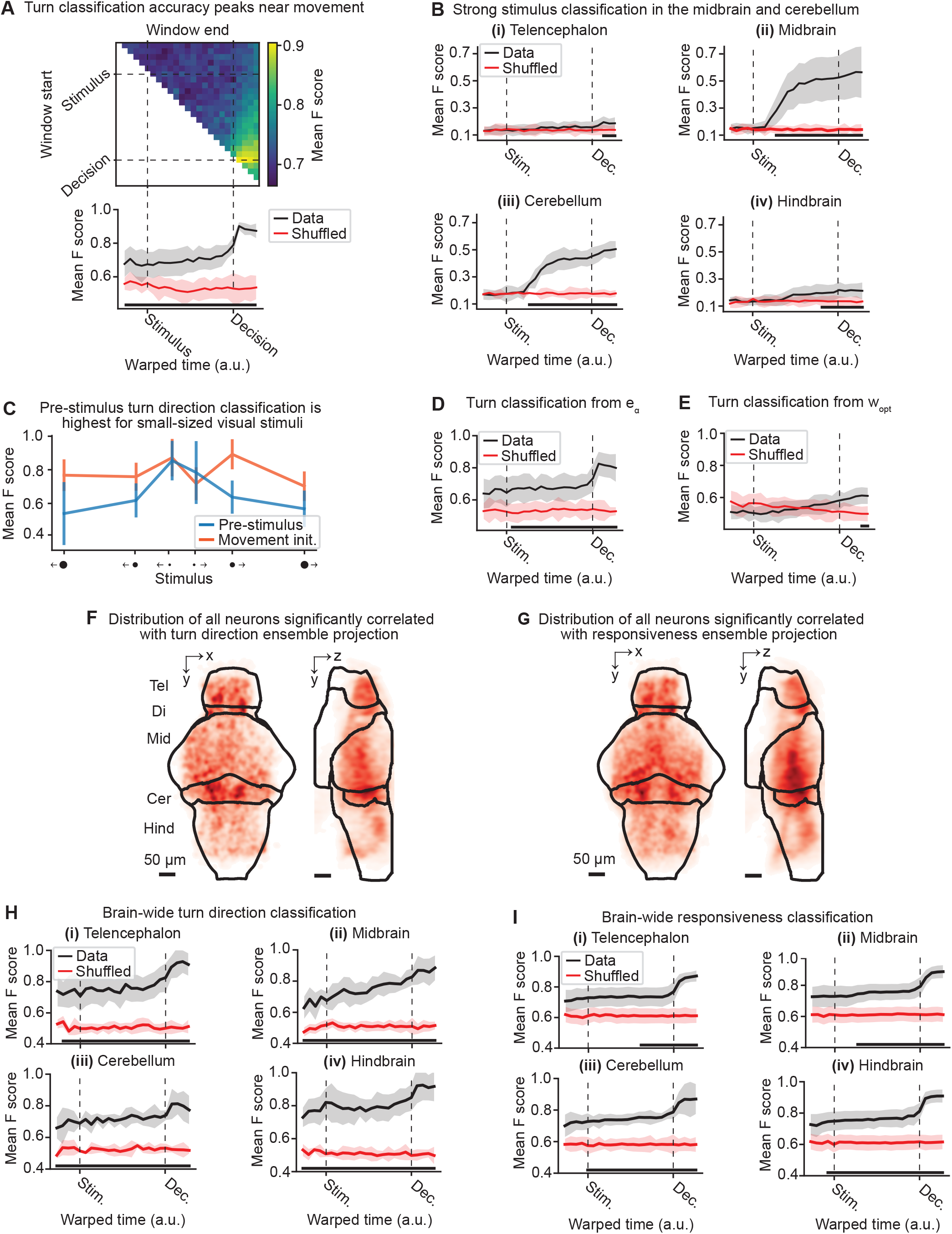
related to Figure 4, pre-motor neuronal populations predictive of single-trial behavior. **A.** Classification of turn direction as a function of time window analyzed. Top: a time window is swept across various start timepoints (y-axis) and end timepoints (x-axis) to determine the classification performance across various time intervals of the neuronal dynamics. Bottom: Mean ± 95% confidence interval of the F score for the best time interval ending at the given timepoint (Data), compared to shuffled data in which the turn direction labels are randomized. The classification accuracy is significantly higher than shuffled data (p<0.05, paired t-test) even before stimulus onset. **B.** Stimulus classification from various brain regions. As in Figure 4B, except only performing classification using neurons in a single brain region. Strong decoding is only observed in the midbrain and cerebellum after stimulus onset. **C.** Turn direction classification as a function of stimulus size. On average before stimulus onset (blue line), the F score is highest for the smallest size stimuli, whereas the F score is relatively constant at the time of movement initiation (orange line, mean ± 95% CI across n=5 larvae). **D.** As in A, except for classifying left or right turn direction from the projections onto the trial-to-trial noise modes 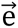_α_, as opposed to the brain-wide dynamics as used in Figure 4D. **E.** As in A, except for classifying left or right turn direction from the projection onto the optimal stimulus decoding dimension 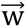_opt_. **F-G.** Distribution of all neurons correlated with the activity of the turn direction bias ensemble (F) or responsiveness ensemble (G). Significant correlations were determined by comparing the observed correlation to that of shuffled datasets where the neuron’s timeseries was randomly circularly permuted. **H-I**. Turn direction (H) and responsiveness (I) exhibit similar predictability across brain regions. As in panels 4D and 4C, respectively, except that classification was performed using neurons in a single brain region.

**Video 1: related to Figure 2, example target-directed and target-avoidance bouts in free behavior.**

On the left, an example target-directed bout showing a 7dpf larval zebrafish approaching a stimulus presented on a screen below the dish (see Figure 2A). On the right, a similar target-avoidance bout from the same larva. The videos are slowed down 10x. These videos correspond to the composite images shown in Figure 2B-C.

## METHODS

### fLFM setup

The sample is illuminated with an LED excitation light (470 nm, Thorlabs M470L4) via Köhler illumination through a standard GFP filter set (Thorlabs MDF-GFP) and a 20×/1.0-NA water immersion objective (Olympus XLUMPLFLN). In order to reduce the amount of excitation light reaching the eyes of the zebrafish, an aluminum mask is placed in the Köhler illumination path conjugate to the sample plane. The imaging path consisted of a f=180 mm Olympus-style tube lens (Thorlabs TTL180-A), a f=180 mm achromatic doublet (Thorlabs AC508-180-A-ML) for the Fourier lens, and a 13×13 mm microlens array with 1.5 mm pitch and r=13.9 mm radius of curvature (OKO Tech APO-GB-P1500-R13.9). The microlens array is mounted inside a five-axis kinematic mount (Thorlabs) to allow precise placement of the array orthogonal to the optical axis. A 5.5-megapixel sCMOS camera (Andor Zyla) is then positioned at the focus of the microlens array to capture the fLFM images.

The theoretical lateral resolution of an fLFM is limited by the numerical aperture *NA*_ML_ of the microlenses and the diffraction-limited spot size at the sensor. For an emission wavelength *λ*, Fourier lens focal length *f*_FL_, objective magnification of *M*, and microlens diameter *d*_MLA_, the Abbe diffraction limit when converted to object space and under the paraxial approximation is given by 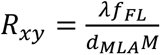. The theoretical axial resolution for a microlens array with a radius *d*_max_ is given by 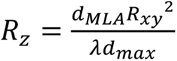 (Guo et al., 2019).

Experimentally, we measured the reconstructed lateral resolution of fLFM as a function of distance from the native image plane (NIP) in Figure S1A using a USAF resolution target and a fluorescent slide. The target was positioned at different axial locations using an automated z-stage, and the corresponding slice of each reconstructed volume was analyzed. An element was considered resolved when its modulation transfer function was greater than 30%. Separately, the reconstructed point-spread function shown in Figure 1C was measured using a 1 µm fluorescent bead embedded in 2.5% agarose, imaged through the water-immersion objective, and positioned at the NIP.

### Zebrafish experiments

For head-fixed zebrafish experiments, n=13 *huc:h2b-gcamp6s* larvae with pan-neuronal and nuclear-localized GCaMP6s (NL-GCaMP6s) expression were imaged 6-9 days post fertilization (dpf). The day before experiments were performed, we immobilized larvae by embedding them in 2.5% agarose approximately 10 mm from the edge of a 35mm petri dish. We then removed the agarose around their eyes and tail to allow for visual stimulation and tail movement. Larvae were not fed after immobilization and were thus in a starved state on the day of the experiment.

The petri dish was covered in a rear projection film (Screen Solutions Int.) such that visual stimuli could be projected directly onto the dish. The projector (BenQ TH671ST) output was filtered with a 610 nm longpass filter (Schott RG610) such that only red light was displayed, removing crosstalk between the stimulation and GCaMP fluorescence. The tail was illuminated with an 850 nm LED and then monitored from below using a near infrared CMOS camera (Ximea MQ013RG-ON).

Visual stimulation was controlled by Stytra (Štih et al., 2019). A short baseline period with no stimulation of 3 to 5 minutes was included at the start of each recording. To check for intact visual circuitry, drifting gratings with a period of 40° (visual angle) and speed of 40°/s to the left or right were shown three times every 30 minutes. Only larvae that displayed a robust optomotor response were further analyzed. The remaining trials consisted of drifting dots that began in the center of the visual field and moved to either the left or right at 30°/s. Dots were shown at maximum contrast against a dark background. Various diameters were shown from 0.3° to 44° visual angle, with 3 to 6 different sizes used in a recording. Each trial lasted 3 seconds and the inter-trial interval was randomly distributed with a mean of 9 seconds and a standard deviation of 3 seconds. This inter-trial interval was chosen empirically to ensure that the visually-evoked activity from the previous trial was negligible during any given trial (see Figure S3C).

### Data acquisition

The fLFM camera was triggered at 5 or 10 Hz using a microcontroller (Adafruit Grand Central). The first trigger was used to initiate the tail behavior camera. The tail was monitored in real-time using Stytra (Štih et al., 2019) at approximately 200 Hz. The recording duration lasted 1 to 2 hours.

### Data processing

Raw fLFM images were denoised using a custom-trained DeepInterpolation (Lecoq et al., 2021) model. DeepInterpolation is a self-supervised approach to denoising, which denoises the data by learning to predict a given frame from a set of frames before and after it. Time-varying signal can be distinguished from shot noise because shot noise is independent across frames, but signal is not. Therefore, only the signal is able to be predicted from adjacent frames. This has been shown to provide a highly effective and efficient denoising method (Lecoq et al., 2021). We trained a “unet_single_1024” model with 5 pre and post frames on a training set of n=20 example recordings and validation set of n=5 recordings. We ran the training until convergence of the validation loss. Denoised raw fLFM images were reconstructed using Richardson–Lucy deconvolution and an experimentally measured PSF obtained from a 1 µm fluorescent bead embedded in 2.5% agarose and sampled through the axial range in 2 µm increments. The Richardson-Lucy algorithm took about 1 second per iteration on a GPU (NVIDIA TITAN V) and we used 12 to 20 iterations per frame. This resulted in a reconstructed volume of approximately 760 × 360 × 280 μm^3^ with voxels of 2.4 μm laterally and 4 μm axially.

Neuronal ROIs and timeseries were then extracted using the CNMF-E (Zhou et al., 2018) variant of the CaImAn software package (Giovannucci et al., 2019), which is optimized for processing of one-photon imaging data. Each plane was analyzed individually and then the results were merged. To determine neuronal ROIs, the SNR threshold was set at 2, the spatial consistency (rval) threshold was set at 0.8, and the CNN-based classifier threshold was set at 0.9. Neuronal timeseries were not deconvolved. Background fluorescence was factored out using the CNMF-E ring model. The planes were then collated by merging any units with a correlation greater than 0.7, lateral distance less than 10 μm, and axial distance less than 16 μm. Overlapping units were merged by summing each timeseries weighted by its SNR. Reported counts are counts of segmented ROIs that met the signal-quality criteria; they should not be interpreted as a complete census of labeled cells, and an ROI cannot always be assumed to correspond to a single neuron. Effects of photobleaching were removed by normalizing by a fit to the mean signal *f*(*t*) across all neurons over the recording with a bi-exponential model *f̂*(*t*) = *ae*^bt^ +*ce*^dt^. Each neuronal timeseries was then z-scored before all following analyses.

For localization of neuronal ensembles to specific brain regions, each recording was aligned to the Z-Brain atlas (Randlett et al., 2015). Alignment was performed by manually marking a number of keypoints on the reference image and the standard deviation projection over time of the 3D fLFM recording, and then estimating the affine transformation between the reference and fLFM volume via least squares.

### Analysis of freely swimming behavior

For freely swimming experiments, n=9 6-9 dpf *huc:h2b-gcamp6s* larvae were individually placed in a 90 mm petri dish with a shallow (∼3 mm) amount of water. The dish was placed above a display (Apple iPad) with a 6 mm piece of clear acrylic in between. Black dots of sizes ranging from 1° to 40° were randomly shown at maximum contrast and moving across the dish in one direction at 30°/s. A camera (FLIR Grasshopper3 GS3-U3-41C6M-C) with a zoom lens (Navitar MVL7000) was mounted above to monitor the larvae’s behavior at >100 Hz.

Each larva’s movements were tracked using DeepLabCut (Mathis et al., 2018), which was trained to track the position of the eyes, the swim bladder, and four points along the spline of the tail. Anipose was used to build a pipeline to process large numbers of videos (Karashchuk et al., 2020). Movement bouts were extracted by applying a threshold to the velocity of the centroid of the tracked points. Only movement bouts in which the centroid was within approximately six body lengths of the visual stimulus were further analyzed. For each bout, we then calculated the mean change in distance to the visual stimulus, with positive values denoting the larvae moving further away from the visual stimulus.

### Analysis of head-fixed behavior

During head-fixed visual experiments, 20 points along the spline of the larva’s tail were tracked in real time using Stytra (Štih et al., 2019). The overall tail curvature was then quantified by summing the angles between each segment of the tail tracking. For comparison with neural data, this was convolved with the NL-GCaMP6s kernel to define the tail direction kernel, while the GCaMP-kernel-convolved absolute tail curvature defined the tail vigor kernel. The angular velocity and angular acceleration were computed for the overall tail curvature summed over all segments. The NL-GCaMP6s kernel was estimated empirically by aligning and averaging a number of calcium events. This kernel corresponds to a half-rise time of 400 ms and half-decay time of 4910 ms. The use of a fixed calcium kernel does not account for any variability in the GCaMP response across cells, which could be due to differences such as cell type or expression level. Therefore, this analysis approach may not capture the full set of neurons which exhibit stimulus correlations but exhibit a different GCaMP response.

Bouts of movement were extracted by thresholding the absolute tail curvature with a value *σ*_active_ set to one standard deviation of the absolute tail curvature over time. To study the behavioral response to various stimuli, the mean tail curvature during active bouts was computed for each stimulus presentation period; trials in which there were no movement bouts were excluded. For turn direction analyses, trials with a mean tail curvature during bouts > *σ*_active_ were considered as left turn trials, whereas trials with a mean tail curvature during bouts < −*σ*_active_ were considered right turn trials.

### Visual stimulus decoding

For the decoding shown in Figure 3B, the stimulus-evoked neural response was taken as the average response of each neuron during a single stimulus presentation. For each larva, a logistic regression model with lasso regularization was trained on all trials using 6-fold cross-validation with 3 repetitions. The lasso regularization parameter, which induces sparsity, was swept and the model with the highest average cross-validated accuracy was used. The confusion matrix, computed on the held-out testing trials, was reported as the average over all rounds of cross-validation.

### Identification of stimulus decoding and noise dimensions

The comparison of the neural dimensions encoding visual stimuli versus trial-to-trial noise was modeled after Rumyantsev et al. (2020). Partial least squares (PLS) regression was used to find a low-dimensional space that optimally predicted the visual stimuli, which we refer to as the visually-evoked neuronal activity patterns. To perform regression, a visual stimulus kernel was constructed by summing the timeseries of each individual stimulus type, weighted by the stimulus size and negated for trials on the right visual field, thus providing a single response variable 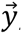 encoding both the location, size, and timing of all the stimulus presentations. This stimulus kernel was then convolved with the temporal response kernel of our calcium indicator (NL-GCaMP6s).

PLS regression identifies the normalized dimensions 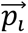 and 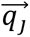 that maximize the covariance between paired observations 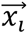 and 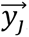, respectively. In our case, the visual stimulus is represented by a single variable 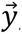, simplifying the problem to identifying the subspace of neural activity that optimally preserves information about the univariate visual stimulus (sometimes referred to as PLS1 regression). That is, the *N* × *T* neural time series matrix *X* is reduced to a *d* × *T* matrix spanned by a set of orthonormal vectors. PLS1 regression is performed as follows:

#### PLS1 algorithm

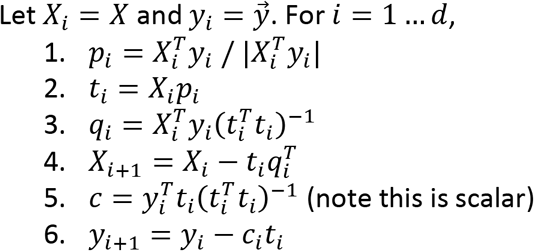

The projections of the neural data {*p*_i_} thus span a subspace that maximally preserves information about the visual stimulus 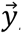. Stacking these projections into the *N* × *d* matrix *P* that represents the transform from the brain-wide neural state space to the visually-evoked subspace, the optimal decoding direction is given by the linear least squares solution 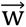_opt_ = *P*(*P*^T^*X*^T^*XP*)^−1^*P*^T^*X*^T^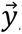. The dimensionality *d* of PLS regression was optimized using 6-fold cross-validation with 3 repeats and choosing the dimensionality between *d* = 1 and 20 with the lowest cross-validated mean squared error for each larva. Then, 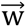_opt_ was computed using all time points.

For each stimulus type, the noise covariance matrix *C* was computed in the low-dimensional PLS space, given that direct estimation of the noise covariances across many thousands of neurons would likely be unreliable. A noise covariance matrix was calculated separately for each stimulus, and then averaged across all stimuli. As before, the mean activity *μ*_i_ for each neuron *i* was computed over each stimulus presentation period. The noise covariance then describes the correlated fluctuations *δ*_i_ around this mean response for each pair of neurons *i* and *j*, where *C*_ij_ = (*δ*_i_ *δ*_j_ ⟩.

The noise modes 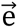_α_ for α = 1 … *d* were subsequently identified by eigendecomposition of the mean noise covariance matrix across all stimuli, 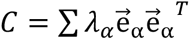. The angle between the optimal stimulus decoding direction and the noise modes is thus given by 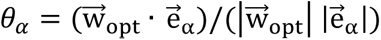.

To assess the reliability of 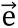_1_ and 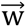_opt_, trials were randomly divided into two non-overlapping halves separately within each stimulus condition. The PLS basis and its dimensionality were defined once for each larva using all available trials, providing a common coordinate system in which directions estimated from the two trial subsets could be compared. Neural activity from each half was projected into this common PLS space. Within each half, 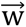_opt_ was estimated independently by least-squares regression of the stimulus variable against the PLS scores. The stimulus-conditioned noise covariance matrices were also calculated using only the trials in that half, averaged across stimulus conditions, and decomposed to obtain the corresponding leading noise direction 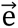_1_. This procedure was repeated for 100 random 50:50 partitions. Because the sign of an eigenvector is arbitrary, alignment was quantified using absolute cosine similarity. We calculated the similarity between 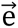_1_ estimated from the two halves, the similarity between 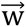_opt_ estimated from the two halves, and the cross-held-out similarities between 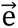_1_ from one half and 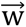_opt_ from the other. The two cross-held-out comparisons were averaged within each partition.

### Classification of single trial turn direction, responsiveness, and stimulus

To predict decisions from neural activity despite the variable reaction times, each trial was time-warped in order to align the stimulus onset and the movement initiation timepoints, following (Lin et al., 2020). Left and right turn trials were extracted as described previously. Response trials included both left and right turn trials (i.e., the absolute value of mean tail curvature > *σ*_active_), whereas nonresponse trials were motionless (absolute mean tail curvature < *σ*_active_). In particular, forward-motion trials were excluded from these analyses. For no response trials, a random reaction time was selected from the responsive trials and the corresponding time-warping was applied. Logistic regression binary classification models with lasso regularization were fit to classify left versus right turn trials or response versus no response trials from various time windows of the time-warped neural activity. Thus, time-warped neuronal activity was averaged over windows from various start and end timepoints to determine the predictability at each time window. The optimal predictability at any given timepoint was then considered as the best time window that ended at that timepoint. For each larva, 6-fold cross-validation with 3 repetitions was utilized to determine the cross-validated F score, accuracy, and confusion matrix. For stimulus classification, logistic regression was similarly applied for multi-class classification using the one-versus-rest scheme. Overall classification performance was quantified using the mean F score across larvae and was compared to shuffled data in which the class labels had been randomly permuted.

Visualizations of single trial trajectories were made by projecting the time-warped trials onto the weights of an example, randomly chosen logistic regression model.

Significant neuronal ROIs contributing to turn direction and responsiveness classification were identified by comparing a neuron’s absolute coefficient across the multiple cross-validation rounds to that from models fit to shuffled class labels. Neurons were deemed to be significantly contributing using a one-tailed t-test between each ROI’s coefficients and its coefficients estimated from shuffled data across the cross-validation rounds, p<0.05. Similarly, neurons were considered to be significantly correlated with the turn direction bias projection and the responsiveness projection in Figures S5F-G using a one-tailed t-test between each neuron’s correlations and its correlations estimated from shuffled data across the cross-validation rounds.

## Notes

### Competing Interest Statement

The authors have declared no competing interest.

### Summary of Updates

This version of the manuscript has been revised to address all comments and issues raised by the reviewers (see authors response letter).

