## Supplementary material for "Whole-brain neural substrates of behavioral variability in the larval zebrafish": Reviewers Response Letter

*Public Reviews:*

*Reviewer #1 (Public review):*

*Summary:*

*In this paper, Manley and Vaziri investigate whole-brain neural activity underlying behavioural variability in zebrafish larvae. They combine whole brain (single cell level) calcium imaging during the presentation of visual stimuli, triggering either approach or avoidance, and carry out whole brain population analyses to identify whole brain population patterns responsible for behavioural variability. They show that similar visual inputs can trigger large variability in behavioural responses. Though visual neurons are also variable across trials, they demonstrate that this neural variability does not degrade population stimulus decodability. Instead, they find that the neural variability across trials is in orthogonal population dimensions to stimulus encoding and is correlated with motor output (e.g. tail vigor). They then show that behavioural variability across trials is largely captured by a brain-wide population state prior to the trial beginning, which biases choice - especially on ambiguous stimulus trials. This study suggests that parts of stimulus-driven behaviour can be captured by brain-wide population states that bias choice, independently of stimulus encoding.*

*Comments on revisions:*

*The authors have revised their manuscript and provided novel analyses and figures, as well as additions to the text based on our reviewer comments.*

*As stated in my first review, the strength of the paper principally resides in the whole brain cellular level imaging - using a novel fourier light field microscopy (Flfm) method - in a well-known but variable behaviour.*

*Many of the authors' answers have provided additional support for their interpretations of results, but the new analysis in Figure 3g - further exploring the orthogonality of e1 and wopt - puts into question the interpretation of a key result: that e1 and wopt are orthogonal in a non-arbitrary way. This needs to be addressed. I have made suggestions below to address this:*

*Reviewer 3 had correctly highlighted the issue that in high-dimensional data, there is an increasingly high chance of two vectors being orthogonal. The authors address this by shuffling the stimulus labels. They then state (and provide a new panel g in Fig. 3) that the shuffled distribution is wider than the actual distribution, and state that a wilcoxon rank-sum test shows this is significant. Given the centrality of this claim, I would like the authors to clarify what exactly is being done here, as it is not clear to me how this conclusion can be drawn from this analysis:*

*In lines 449:453 the authors state:*
*'While it is possible to observe shuffled vectors which are nearly orthogonal to e1, the shuffled distribution spans a significantly greater range of angles than the observed data (p<0.05, Wilcoxon rank- sum test), demonstrating that this orthogonality is not simply a consequence of analyzing multi-dimensional activity patterns. '*
*I don't understand how the authors arrive at the p-value using a rank-sum test here. (a) What is the n in this test? Is n the number of shuffles? If so, this violates the assumptions of the test (as n must be the number of independent samples and not the arbitrary number of shuffles). (b) If the shuffling was done once for each animal and compared with actual data with a rank-sum test, how likely is that shuffling result to happen in 10000 shuffle comparisons?*
*I am highlighting this, as it looks from Figure 3g that the shuffled distribution is substantially overlapping with the actual data (i.e., not outside of the 95 percentile of the shuffled distribution), which would suggest that the angle found between e1 and wept could happen by chance.*

*I would also suggest the authors instead test whether e1 is consistently aligned with itself when calculated on separate held out data-sets (for example by bootstrapping 50-50 splits of the data). If they can show that there is a close alignment between independently calculated e1's across separate data sets (and do the same for wopt), and then show e1 and wopt are orthogonal, then that supports their statement that e1 and wopt are orthogonal in a meaningful way. Given that e1 captures tail vigor variability (and Wopt appears to not) then I would think this could be the case. But the current answer the authors have given is not supporting their statement.*

We thank the reviewer for their recognition of the strength of our imaging approach, as well as for raising the concerns regarding the analysis shown in the previous version of Figure 3G. In the following we are providing responses to each of the specific points raised by the reviewer. We realize that our description of the statistical approach was not sufficiently clear. To answer the reviewer's questions: (a) n was not the number of arbitrary shuffles but indeed the independent biological sample size which was n=8 larvae. Further, the shuffled realizations (10,000 per animal) were intended to characterize a null distribution and were not intended to substitute the number of independent biological samples. We agree with the reviewer that the rank-sum comparison as previously presented did not directly test how likely the observed alignment was to arise by chance under the shuffled null distribution, which is the question raised by the reviewer in part (b). Rather than retaining or further interpreting this comparison, we have removed the shuffled analysis, its associated p-value, and the statement that it demonstrated that the observed orthogonality could not arise from the dimensionality of the activity space and have instead adopted the reviewer's suggested reliability analysis.

Following their suggestion of 50:50 splits, we interpreted "separate held-out datasets" as non-overlapping sets of trials within each larva's recording. Vectors cannot be compared directly across recordings because each recording contains a distinct neuronal state space. For each larva, we repeatedly divided the trials for every stimulus condition into two non-overlapping halves. To express the directions estimated from both halves in a common coordinate system, the PLS basis was defined once for each larva's recording. Within the common basis, $\vec{e}_1$ was then estimated from the stimulus-conditioned noise covariance in each half separately, and $\vec{w}_{opt}$ was similarly estimated in each half separately by regression against the stimulus variable.

We then quantified alignment using absolute cosine similarity because the sign of an eigenvector is arbitrary. New Figure 3G shows the similarity between two separately estimated $\vec{e}_1$ vectors ("split-half"), two separately estimated $\vec{w}_{opt}$ vectors ("split-half"), and the cross-held-out similarity between $\vec{e}_1$ estimated from one half and $\vec{w}_{opt}$ estimated from the other ("opposite-half"). Each point represents the median across repeated splits for one larva, with whiskers showing the within-larva interquartile range.

Both population directions were reproducible across trial subsets: the median of the per-larva split-half similarities was approximately 0.88 for $\vec{e}_1$ and 1.0 for $\vec{w}_{opt}$. In contrast, the median cross-held-out similarity between $\vec{e}_1$ and $\vec{w}_{opt}$ was approximately 0.003. Thus, $\vec{e}_1$ and $\vec{w}_{opt}$ are individually stable directions, yet remain nearly orthogonal when estimated from non-overlapping trials. This directly implements the criterion proposed by the reviewer and supports our interpretation that the observed geometry reflects two reproducible and distinct population directions, rather than instability in either estimate. We have revised the Results, Methods, and Figure 3 legend accordingly.

We thank the reviewer for their valuable suggestion to highlight this analysis, which we believe will make the reliability and orthogonality of our population direction estimates clearer and more quantitative for the reader.

*Reviewer #2 (Public review):*

*This work by Manley and Vaziri identify brain networks that are associated with trial-to-trial variability during prey-capture and predator avoidance behaviors. However, mixing of signals across space and time make it difficult to interpret the data generated and relate the data to findings from prior work.*

*Comments on revisions:*

*In their response to prior reviewer comments, Manley and Vaziri have now provided helpful methodological clarity and additional analyses. The additional work makes clear that the claims of variability and mixing of sensory, motor, and internal variables at the single-cell level are not well supported.*

*RESOLUTION*

*- The new information provided regarding resolution may not be very relevant as this was from an experiment in air. It would be much more informative to show how PSF degrades in the brain with depth.*

We thank the reviewer for emphasizing this distinction. We agree that measuring the depth-dependent PSF within the brain would be more directly relevant to *in vivo* performance. However, we would like to clarify that the shown measurement in Figure 1C was not performed in air, as the reviewer suggests. We characterized the optical resolution using two complementary approaches. First, we measured the modulation transfer function using a USAF resolution target, providing a characterization of the spatial frequencies resolved by the imaging system (Figure S1A). Second, we measured fluorescent beads embedded in 2.5% agarose, providing a PSF measurement in a more tissue-relevant optical environment (Figure 1C). Because a USAF target cannot practically be embedded within the zebrafish brain, these complementary measurements provide a feasible characterization of system resolution under the experimental constraints.

To make this distinction more explicit and clearly distinguish between the two different approaches of PSF measurements, we have revised the Methods section of our revised manuscript as follows: "*Experimentally, we measured the reconstructed lateral resolution of fLFM as a function of distance from the native image plane (NIP) in Figure S1A using a USAF resolution target and a fluorescent slide. The target was positioned at different axial locations using an automated z-stage, and the corresponding slice of each reconstructed volume was analyzed. An element was considered resolved when its modulation transfer function was greater than 30%. Separately, the reconstructed point-spread function shown in Figure 1C was measured using a 1 μm fluorescent bead embedded in 2.5% agarose, imaged through the water-immersion objective, and positioned at the NIP.*"

We also extended the figure legends to identify the Figure 1C sample as *"a 1 μm fluorescent bead embedded in 2.5% agarose"* and to describe Figure S1A as *"reconstructed lateral resolution of fLFM and cLFM across the axial imaging range, measured using a USAF resolution target translated relative to the native image plane."*

However, on a broader note, we acknowledge that neither measurement fully captures the depth dependence introduced by real biological tissue. Directly measuring the fLFM detection PSF at defined locations within a living brain would require generating a spatially confined fluorescent source at multiple depths (for example, using a hybrid two-photon excitation/fLFM instrument) which is not available for the present system and would require a substantially different experimental setup. To our knowledge, a direct depth-dependent measurement of the LFM detection PSF within the intact larval zebrafish brain has not previously been reported. Consistent with established practice in the field, LFM systems are generally characterized using fluorescent beads or resolution targets in controlled calibration preparations, including Cohen et al. (2014), Cong et al. (2017), and Quicke et al. (2020). These standardized measurements provide a reproducible characterization of system performance across imaging configurations.

Published work nevertheless provides relevant context for the likely magnitude of tissue-induced focal broadening in the larval zebrafish brain. Favre-Bulle et al. (2015) combined *in vivo* measurements of backscattered 488 nm light with Monte Carlo modeling of the focal intensity distribution in the larval zebrafish brain. In their model, the focal FWHM in the absence of scattering was 0.13 μm laterally and 0.29 μm axially; after propagation to a depth of 100 μm in zebrafish brain tissue, this increased to 0.16 μm laterally and 0.39 μm axially. They concluded that single-neuron-sized illumination volumes are achievable throughout the entire brain at this wavelength. The very low refractive index contrast driving scattering in this model (n = 1.35 for nuclei versus n = 1.34 for surrounding tissue) confirms that the larval zebrafish brain is an inherently low-scattering preparation. Although this analysis concerns illumination rather than the fLFM detection PSF and therefore does not provide a direct comparison with our agarose measurements, it suggests that tissue-induced broadening at comparable visible wavelengths may be relatively modest over the larval zebrafish brain. Accordingly, we have revised the Results section of our revision stating: *“These measurements provide a standardized characterization of fLFM optical performance under controlled calibration conditions. Previous work combining in vivo backscatter measurements with experimentally constrained modeling at a comparable visible wavelength found relatively modest focal broadening over the first 100 μm of larval zebrafish brain, including within nucleus-rich regions (Favre-Bulle et al., 2015). Nevertheless, cellular separability in vivo may vary with imaging depth, labeling density, signal quality, and reconstruction.”*

Separately, we note that Nöbauer et al. (2017) benchmarked LFM source localization and activity extraction against simultaneous two-photon recordings in the more strongly scattering mouse brain. We cite this as evidence that computational source-extraction methods applied to LFM measurements can localize neuronal sources and recover their activity under conditions of substantially greater tissue scattering than those encountered in the larval zebrafish brain. Taken together, we believe that the PSF and the location precision of our system were sufficient to resolve individual neurons over much of the imaged depth range in the semitransparent zebrafish brain.

*DEPTH*

*- It is helpful to see the registered light-field and confocal images. Both appear to provide poor or little information in regions >200 below the surface (like the hypothalamus), making the claim that whole brain data is being collected at cellular resolution difficult to justify.*

We thank the reviewer for raising this important distinction. We agree that the reconstructed volume encompasses the gross anatomical extent of the larval zebrafish brain, but that labeling, signal quality, and effective neuronal sampling are not uniform across all depths. In particular, deep ventral regions, including the hypothalamus, may be underrepresented. We have therefore revised the text to distinguish between anatomical coverage and uniform cellular-resolution sampling: "*Although the larval zebrafish brain is relatively transparent and only weakly scattering, which should mitigate depth-dependent degradation relative to more opaque vertebrate preparations, in vivo cellular separability at depth can still vary with reporter brightness, neuronal density, and residual scattering*."

We emphasize, however, that our original use of "whole-brain imaging" is consistent with the generally established terminology in the zebrafish literature: The *huc:h2b-rfp* reference image shown in Figure S1E was reproduced from Randlett et al. (2015), titled *"Whole-brain activity mapping onto a zebrafish brain atlas."* The same pan-neuronal nuclear-labeling approach and the *huc:h2b-gcamp6s* line used in our functional experiments have repeatedly been described across literature as enabling *"whole-brain imaging,"* including in Vladimirov et al. (2014), Migault et al. (2018), Bruzzone et al. (2021), de Vito et al. (2022), and van der Plas et al. (2023).

Nevertheless, to avoid implying uniform labeling, neuronal sampling, or cellular separability across every anatomical region, we have revised the manuscript to distinguish explicitly between whole-brain anatomical coverage and the brain-wide distribution of the sampled neuronal population. We retain the term "whole-brain imaging" in its established zebrafish-literature sense, while qualifying statements that could otherwise be interpreted as implying uniform cellular-resolution sampling throughout the brain. In addition, we now distinguish the optical resolution measured with fluorescent beads or a resolution target from effective cellular resolution *in vivo.* The revised Results state: "*Although the larval zebrafish brain is relatively transparent and only weakly scattering, which should mitigate depth-dependent degradation relative to more opaque vertebrate preparations, in vivo cellular separability at depth can still vary with reporter brightness, neuronal density, and residual scattering."*

Figure S1E shows that the depth-dependent reduction in detected ROIs broadly resembles the labeling distribution in the Z-Brain reference image. This comparison indicates that reporter labeling contributes to the reduced representation of deep regions such as the hypothalamus and that this pattern is not unique to our fLFM recordings. However, we agree that the comparison cannot establish that fLFM contributes no additional depth-dependent limitation. We have therefore revised the caption of Figure S1E to state: *"The distribution of neuronal ROIs in the fLFM recordings is broadly similar to the reference image across major brain divisions; however, labeling and detected ROI density are reduced in deep ventral regions, including the hypothalamus. Improved pan-neuronal transgenic lines or more targeted labeling may be desired for these regions."*

We believe these revisions preserve the established meaning of "whole-brain imaging" in the zebrafish literature while adopting more conservative terminology for our biological conclusions and explicitly acknowledging the nonuniform cellular sampling of deep brain regions.

*MERGING*
*- The typical soma at these ages has a radius of 2.5 microns, which corresponds to a volume of 65 microns^3. Given the close packing of most cells, this means that a typical ROI of 750 microns^3 contains more than 10 neurons. Therefore, the authors should not claim they are reporting activity at cellular resolution.*

*- Furthermore, the fact that these ROIs contains tens of cells brings into question the degree of variability at the single-cell level. For example, if every cluster of 10 cells has one variable cell, then all clusters might be labeled as exhibiting variability even though only 10% of the cells show variability*

As described above, we have updated our main text to make it clear that the size and density of neurons vary greatly across the larval zebrafish brain, which has an impact on the ability to resolve individual neurons. The most important consideration in LFM is the interaction between cell density in the specimen and the depth-dependence of the microscope's PSF. For this reason, during experiments, we placed the native image plane of the microscope approximately in the center of the midbrain, and particularly the optic tectum. This ensured that our resolution was highest in a region with dense cell bodies that was crucial to the analyses presented in Figure 3.

We would also like to clarify that the size and distribution of ROIs are not always strong indicators of optical resolution. In particular, we used CNMF-E for ROI segmentation, which essentially identifies groups of adjacent voxels that are highly correlated. For these reasons, highly correlated neighboring neurons may also appear to be joined in a single ROI. Still, we have thoroughly revised the text of our manuscript, qualifying our resolution capabilities as dependent on the brain region, cell type, and depth of imaging. Additionally, we refrain from making any claims about the absolute number of individual neurons that exhibit variability. Throughout the revised text, we have referred to the individual units as ROIs, rather than neurons; for example: "*Across our 1-2 hour recordings, we identified 16,524 $\pm$ 3,942 neuronal ROIs per animal*".

*SLOW CALCIUM DYNAMICS*

*- Convolution/Deconvolution with the inappropriate kernel both have problems, some of which the authors have noted. However, by not deconvolving, the authors are significantly obscuring the interpretation of their data by mixing together signals across time.*

We thank the reviewer for highlighting this point. We agree that the slow kinetics of nuclear-localized GCaMP6s mix fluorescence signals across time and that it is important to determine whether this temporal filtering affects our conclusions. For the primary analyses, we analyzed the extracted fluorescence signals rather than inferred spike trains because deconvolution necessarily depends on assumptions about the calcium-response kernel. In particular, the response kinetics of *huc:h2b-gcamp6s* may vary across neurons because of differences in cell type, indicator expression, signal-to-noise ratio, and other biological or measurement factors, which the reviewer has pointed out in their previous arguments. In the absence of simultaneously recorded electrophysiological ground truth, deconvolution cannot be assumed to remove this temporal mixing completely. Therefore, we designed our study to account for the long-timescale decay of GCaMP and the highly directional nature of such "mixing", as we have previously described.

Nevertheless, we agree that deconvolution can provide a complementary sensitivity analysis. Following the reviewer's suggestion, we therefore repeated the key analyses underlying Figures 3 and 4 using neuronal time series deconvolved with constrained FOOPSI (Vogelstein et al., 2010; Pnevmatikakis et al., 2016). Constrained FOOPSI was applied independently to each neuron's CNMF-E fluorescence time series using a first-order autoregressive model and the OASIS solver. This allowed the decay parameter to be estimated separately for each neuron. We retained the outputs without thresholding or binarization. We excluded failed or non-finite results, traces with an invalid autoregressive coefficient, constant fluorescence traces, and estimates containing fewer than 20 positive time bins. We refer to these values as continuous-valued deconvolved activity estimates rather than inferred spikes because their relationship to the underlying spike trains cannot be established without ground-truth electrophysiology.

We present these analyses in Response Figure 1. As expected, the visually evoked response was substantially sharper and decayed more rapidly after deconvolution

(Response Figure 1A), demonstrating that constrained FOOPSI reduced the long fluorescence decay present in the original signals.

Importantly, the principal population-level conclusions were preserved. When the signal/noise geometry analysis from Figure 3E was repeated using the deconvolved activity estimates, the dominant noise modes remained nearly orthogonal to the optimal visual-stimulus decoding direction (Response Figure 1B). A paired comparison showed that the leading noise mode remained nearly orthogonal to the visual decoding direction after deconvolution, although the deconvolved estimates were more variable across larvae (Response Figure 1C). This increased spread may reflect the sensitivity of deconvolution to measurement noise, sparse activity, and uncertainty in the neuron-specific calcium-decay estimates. Importantly, the deconvolved angles remained concentrated near 90 degrees.

We also repeated the time-resolved behavioral decoding analyses from Figures 4C–D using the deconvolved activity estimates. Responsiveness remained decodable before stimulus onset and became increasingly predictable near movement initiation (Response Figure 1D). Turn direction showed a smaller but still above-null pre-stimulus difference, and decoding increased substantially toward movement initiation (Response Figure 1E). Thus, the same qualitative temporal structure observed in the original analysis was retained.

Importantly, we do not interpret the FOOPSI estimates as ground-truth neural firing, nor do we claim that deconvolution completely eliminates temporal mixing. Rather, this complementary analysis demonstrates that reducing the slow calcium decay does not change our principal conclusions: the dominant trial-to-trial variability remains largely orthogonal to the visual decoding dimension, and population activity remains predictive of single-trial responsiveness and turn direction before stimulus onset and increasingly so near movement initiation.

**Response Figure 1**: related to Figures 3 & 4. Key population-level conclusions are preserved after constrained FOOPSI deconvolution.

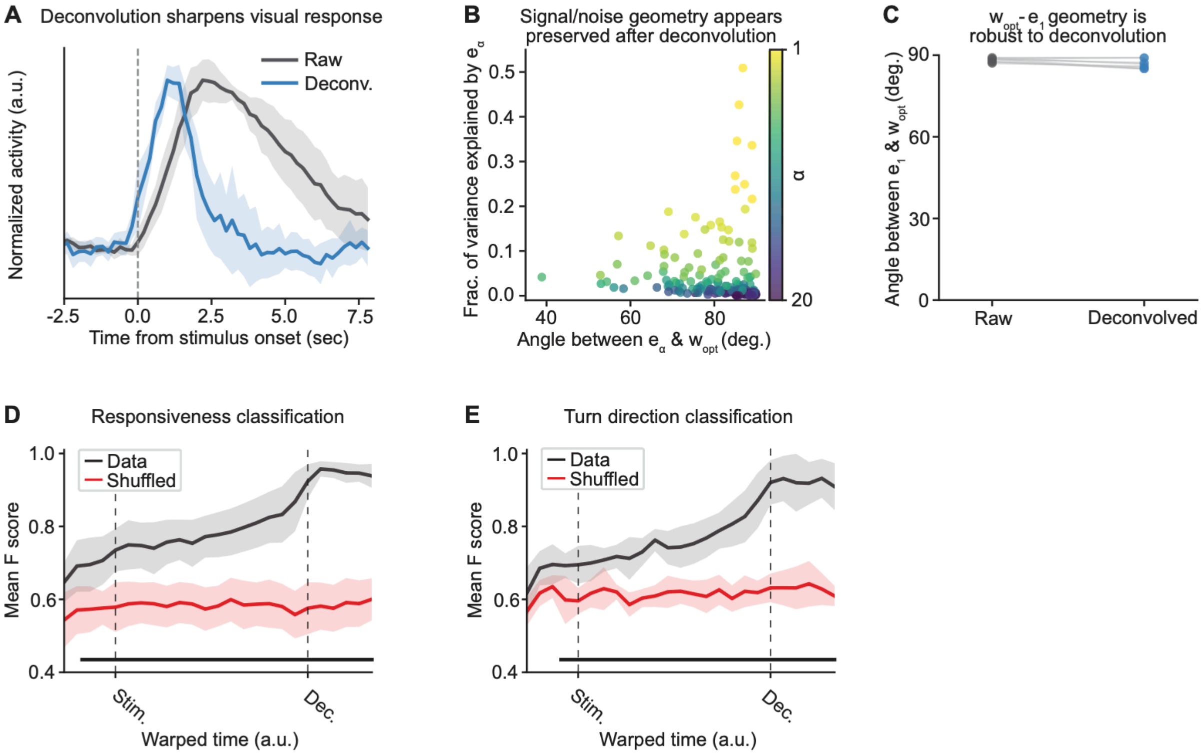


**Response Figure 1: related to Figures 3 & 4. Key population-level conclusions are preserved after constrained FOOPSI deconvolution.**

**A.** Deconvolution shortens the visually evoked response. Mean visually evoked activity aligned to stimulus onset is shown for the original CNMF-E fluorescence signal ("Raw", black) and the FOOPSI-deconvolved activity estimate (blue) across n=8 larvae. Each representation was baseline-centered and normalized independently; therefore, this panel compares response kinetics rather than absolute signal magnitude. Lines indicate the mean across larvae and shaded regions indicate the 95% confidence interval. The dashed vertical line denotes stimulus onset. The FOOPSI-deconvolved response rises and decays more rapidly than the original fluorescence signal.

**B.** Signal/noise geometry is preserved in the FOOPSI-deconvolved activity. The signal/noise geometry analysis in Figure 3E was repeated using the continuous-valued deconvolved activity estimates from n=8 larvae. Each point represents one noise mode $\vec{e}_{\alpha}$. The fraction of trial-to-trial variance explained by each mode is plotted against its angle relative to the optimal visual-stimulus decoding direction ($\vec{w}_{opt}$). Color indicates noise-mode rank ($\alpha$), ordered by decreasing variance explained. The dominant, high-variance noise modes remain close to orthogonal to the visual decoding direction, whereas the lower-variance modes exhibit a broader distribution of angles.

**C.** The leading signal/noise angle is similar before and after deconvolution. Paired comparison of the angle between $\vec{w}_{opt}$ and the leading noise mode $\vec{e}_1$ estimated from the original fluorescence signal (“raw”, black) and from the FOOPSI-deconvolved activity (blue). Each connected pair represents one larva (n=8). The leading noise mode remains nearly orthogonal to the visual decoding direction after deconvolution.

**D-E.** Responsiveness and turn direction, respectively, remain predictable from deconvolved population activity. The Figure 4C-D time-resolved classification analysis was repeated using the FOOPSI-deconvolved activity estimates from n=7 larvae. At each warped time point, the black curve shows the mean cross-validated F score. The red curve shows a shuffled control in which class labels were permuted separately within each stimulus side and size. Lines indicate the mean across larvae and shaded regions indicate the 95% confidence interval. Dashed vertical lines denote stimulus onset (Stim.) and the movement-initiation/decision time point (Dec.). Responsiveness and turn direction are above the shuffled control before stimulus onset and become increasingly predictable near movement initiation.

*- Also, the claim that "neurons highly tuned to a particular stimulus exhibited variability in their responses across multiple presentations of the same stimuli" should be clarified or qualified. It is not clear from what has been shown if the responses are indeed variable, or rather if there is additional activity (or apparent activity) occasionally present that shifts the pre-stimulus baseline around (for example, 3J suggests that in many cases the visual signal from the prior trial is still present when a new trial begins).*

We thank the reviewer for raising this important distinction. Trial-to-trial differences in the activity of visually-tuned neurons can arise either from differences already present at stimulus onset (i.e., shifts in pre-stimulus state) or from differences in the activity trajectory after stimulus onset. The former would shift the apparent baseline of the raw traces, whereas the latter would remain after alignment of each trace to its activity at stimulus onset. We have clarified this distinction in the revised Results text: “*Alternatively, the baseline pre-stimulus state of visually-evoked neurons could be shifted by other circuit components”.*

Both forms of variation are relevant to our results. Figures 4E-G show that pre-stimulus activity patterns differ across behavioral outcomes and ultimately are predictive of turn direction. In revised Figure 3A (discussed below), each trace is offset-corrected to its value at stimulus onset; substantial differences in post-stimulus trajectories remain visible. Thus, the variability shown in Figure 3A is *not* explained by trial-to-trial shifts in activity at stimulus onset.

We also considered whether residual activity elicited by the preceding trial could contribute to these differences. Although this possibility cannot be completely excluded, the average visually-evoked activity has returned, or nearly returned, to baseline by the beginning of the subsequent trial (Figure S3C). More importantly, as we demonstrated in previous versions of the manuscript, the visual component $\vec{w}_{opt}$ is *not* predictive of the observed behavioral variability (Figure S5E), whereas the noise component does. Given the orthogonality of the noise component and the visual component, we find no evidence that residual visually-evoked activity from the preceding trial accounts for the ability to predict behavioral variability.

*- Figure 3A should show when the stimulus occurs, should show some of the prestimulus period, and ideally be off-set corrected so all traces in a given panel start at the same y-value at the beginning of the stimulus period.*

We appreciate the reviewer's suggestions. Following their suggestion we have updated Figure 3A. The figure now includes the pre-stimulus period and explicitly marks the 3-s visual stimulus period. We also offset-corrected each trace so that all trials begin at the same value at stimulus onset. Although pre-stimulus activity is biologically relevant (in fact, Figures 4E-G show that *pre-stimulus* brain state predicts turn direction), this normalization demonstrates that substantial trial-to-trial variability remains in the visually evoked dynamics after controlling for differences in activity at stimulus onset.

*ORTHOGONALITY*

*- It is now clearer that the visual signal and noise vectors were determined for the entire time series with all trials. Therefore, the concern that sources of activation in advance of a given trial were being ignored is alleviated. The concern remains, however, that these sources are being properly accounted for given potential kernel variations and nonlinearity. Nonetheless, it is recognized that the GCaMP filtering most likely would lead to a decrease in the disparity between two populations.*

We thank the reviewer for acknowledging that computing the visual signal and noise vectors across the full time-series helps account for activity preceding individual trials. We agree that variation in GCaMP kinetics and fluorescence nonlinearity could still affect this transformation and cannot be fully eliminated. However, as described above, our complementary analyses using deconvolved activity yielded consistent results, providing additional support that the observed population-level disparity is not driven solely by GCaMP filtering.

*- The authors' clarification that the analyzed ROIs consist of cell clusters raises the trivial possibility that the observed orthogonality between the visual signal and leading noise vectors is explained by noise simply reflecting the activation of different motor or*

*motor-planning related neurons in an ROI, neurons that are separate from visually-encoding neurons in the same cluster.*

We agree with the reviewer that, because some ROIs could contain small clusters of neurons, we cannot exclude the possibility that distinct neurons within an ROI preferentially contributed to visually-evoked or motor-related activity. However, any potential cellular segregation would not itself explain the orthogonality reported here. Both $\vec{w}_{opt}$and $\vec{e}_1$ are defined in the state space spanned by the activity of the measured ROIs. Orthogonality therefore indicates that the visually-evoked response and dominant trial-to-trial variability have distinct patterns of loading across ROIs. If different cellular subpopulations contribute to these components within a given ROI, their activity would be combined in the same measurement coordinate. Such sub-ROI segregation would not be expected to trivially produce orthogonality in the measured ROI state space. On the contrary, pooling visual and motor-related neurons within the same ROI would generally mix their contributions into shared measurement coordinates and reduce the observed orthogonality. The measured orthogonality therefore requires that their aggregate contributions differ systematically across ROIs.

We have revised the Results text to clarify that our result demonstrates separation between population-level coactivation patterns at the spatial resolution provided by fLFM: "*Given that our data do not establish that every ROI corresponds to a single neuron, the observed orthogonality demonstrates separation between population-level coactivation patterns at the spatial resolution provided by fLFM, without excluding the possibility of functional heterogeneity within some ROIs*."

*SOURCES of VARIABILITY*
*- The data presented in Supplemental Figure 3Ei actually is suggestive that eye movements are a significant contributor to the reported variability. Notice how in (1 4) vs (1 5) and (4 7) vs (4 8) there is a notable difference in the distribution of responses. Adding eye kinematic variables to the analysis of Figure S4 could be clarifying.*

We agree with the reviewer's implication that the analyses presented in Figure S3E(i) do not rule out eye movements as one source of neuronal variability. Rather, our intention was to demonstrate that neuronal variability is present even when accounting for some of the eye position and kinematics. Given the multi-dimensional nature of the observed noise correlations (Figures 3D-E), we hypothesize there are multiple sources of variability, as we described in our previous version of the manuscript: *"while movements of the eye likely contribute to aspects of the observed neural variability, they do not dominate the observed neural variability."* In our revised manuscript, we removed the anecdotal claim that "*the magnitude and temporal dynamics of the stimulus-evoked responses show apparently similar levels of variability",* given that we have not shown careful quantification of these features. We look forward to future efforts which can utilize our data to carefully investigate the relationship between neuronal variability, eye kinematics, and behavior.

In response to the reviewer's suggestion, we have updated Figure S4A to include eye kinematic variables. In particular, the angular velocity, and absolute angular velocity of each eye have been included in the correlation analyses. New Figure S4A(ii) demonstrates that the first ten noise modes exhibit varying degrees of correlation with these eye variables. Still, the tail vigor and curvature-related variables exhibit the strongest correlations with these top noise modes. This does not, however, exclude the possibility that additional noise modes (or aspects of eye kinematics not captured by these measures) are more strongly related to eye movements.
